# Insights into gene conversion and crossing-over processes from long-read sequencing of human, chimpanzee and gorilla testes and sperm

**DOI:** 10.1101/2024.07.05.601967

**Authors:** Peter Soerud Porsborg, Anders Poulsen Charmouh, Vinod Kumar Singh, Sofia Boeg Winge, Christina Hvilsom, Marta Pelizzola, Sandra Laurentino, Nina Neuhaus, Asger Hobolth, Thomas Bataillon, Kristian Almstrup, Søren Besenbacher, Mikkel Heide Schierup

## Abstract

Homologous recombination rearranges genetic information during meiosis to generate new combinations of variants. Recombination also causes new mutations, affects the GC content of the genome and reduces selective interference. Here, we use HiFi long-read sequencing to directly detect crossover and gene conversion events from switches between the two haplotypes along single HiFi-reads from testis tissue of humans, chimpanzees and gorillas as well as human sperm samples. Furthermore, based on DNA methylation calls, we classify the cellular origin of reads to either somatic or germline cells in the testis tissue. We identify 1692 crossovers and 1032 gene conversions in nine samples and investigate their chromosomal distribution. Crossovers are more telomeric and correlate better with recombination maps than gene conversions. We show a strong concordance between a human double-strand break map and the human samples, but not for the other species, supporting different PRDM9-programmed double-strand break loci. We estimate the average gene conversion tract lengths to be similar and very short in all three species (means 40-100 bp, fitted well by a geometric distribution) and that 95-98% of non-crossover events do not involve tracts intersecting with polymorphism and are therefore not detectable. Finally, we detect a GC bias in the gene conversion of both single and multiple SNVs and show that the GC-biased gene conversion affects SNVs flanking crossover events. This implies that gene conversion events associated with crossover events are much longer (estimated above 500 bp) than those associated with non-crossover events. Highly accurate long-read sequencing combined with the classification of reads to specific cell types provides a new, powerful way to make individual, detailed maps of gene conversion and crossovers for any species.

## Introduction

Meiotic recombination is a ubiquitous cellular process, and the pathways involved are ultra- conserved. Yet recombination rates can readily be subjected to selection, and recombination hotspots evolve rapidly both within genomes and across species. To gain a better understanding of how and why recombination evolves, we require a deeper mechanistic understanding of the pathways underpinning it and the ability to quantify differences in recombination rates and patterns at the individual level.

In most species, proper meiotic chromosome segregation relies on double-strand breaks (DSBs), where a minority of breaks are initially resolved as crossovers (COs) and the majority are subsequently repaired by the homologous strand leading to non-crossovers (NCOs). In humans, a few hundred DSBs are formed in each meiosis at specific hotspots associated with the presence of PRDM9 motifs^1–3^. In males, DSBs are enriched at the telomeres with further telomeric enrichment in those resolved as COs ^4^. DSBs are repaired by strand invasion from the homologous chromosome, and this process increases the occurrence of *de novo* mutations^5^. In case of a heterozygous position, a mismatch will be induced, which is subsequently repaired by mismatch repair (MMR). If repaired by the incoming strand, this will cause a visible NCO, also known as a gene conversion (GCV). When the heterozygous position contains a weak (A,T) and a strong (C,G) allele, the GCV process is biased towards the strong allele ^6,7^. Such GC-biased gene conversion (gBGC) is powerful in shaping genome evolution. It causes the equilibrium GC frequency to be higher in regions with high recombination rates and leads to a higher GC content genome-wide than the equilibrium expected from mutational processes alone ^8,9^.

Human CO patterns have been extensively studied from linkage disequilibrium (LD) patterns^10^ and pedigrees, with large differences in rate and positions observed between sexes ^11^, and also among individuals of the same species^12^. Comparatively, less is known about GCVs from the resolution of NCOs since these are very hard to detect from LD patterns and require very accurate genotyping to detect from pedigrees^13^. Furthermore, the number of events in each trio is limited, thus making it hard to quantify interindividual variation in both COs and GCVs associated with NCOs.

Highly accurate long-read sequencing offers a new avenue for the detection of both COs and GCVs. Since reads are sufficiently long (typically 10-20 kb), transfer between the paternal and maternal haplotypes will be displayed directly in the read (Figure 1A). In that context, both sperm samples and testicular tissue contain a substantial number of cells that have undergone meiosis and thus are directly informative about COs and GCVs if sequenced. Here, we use high-coverage HiFi sequencing to directly detect meiotic recombinations (COs and GCVs) in sperm and testis samples from humans, chimpanzees, and western gorillas. We report similar short GCV tracts across samples and species but distinct genomic locations of COs and GCVs.

**Figure 1:**
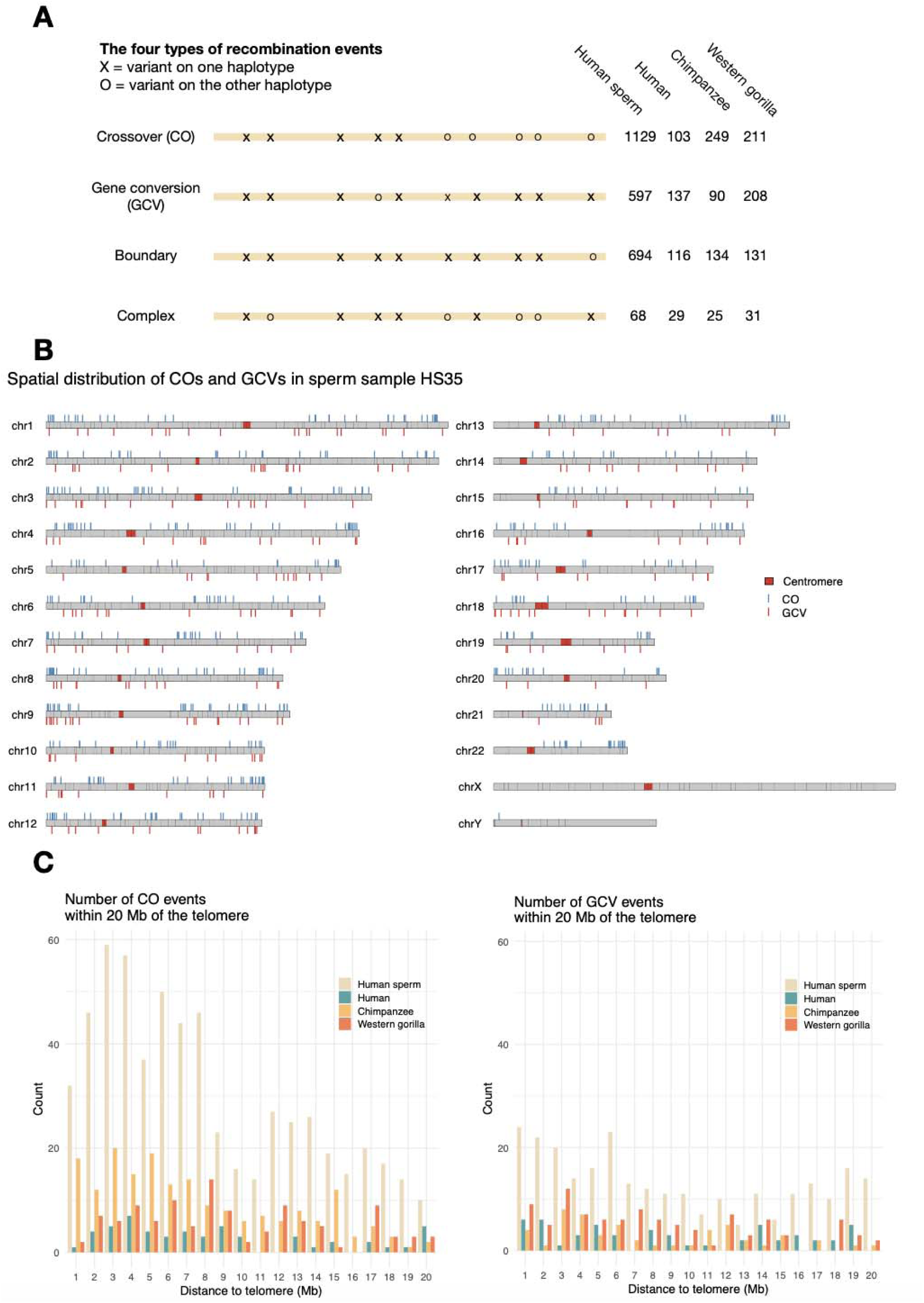
Physical placement of recombination events. **A.** Polymorphism patterns indicating the different types of recombinations identified. O and X marks variants at the two different haplotypes. The total number of each type of event pooled by species and tissue type is shown to the right. **B.** The genomic placement (on the T2T-CHM13v2.0 assembly) of each of the identified CO (above the chromosome) and GCV (below) events for the human sperm sample HS35 (see Table 1). Other samples are shown in the supplement. **C.** The number of crossovers and gene conversions in the 20 Mbp regions closest to the telomere for each tissue type and species separately.

**Table 1:**
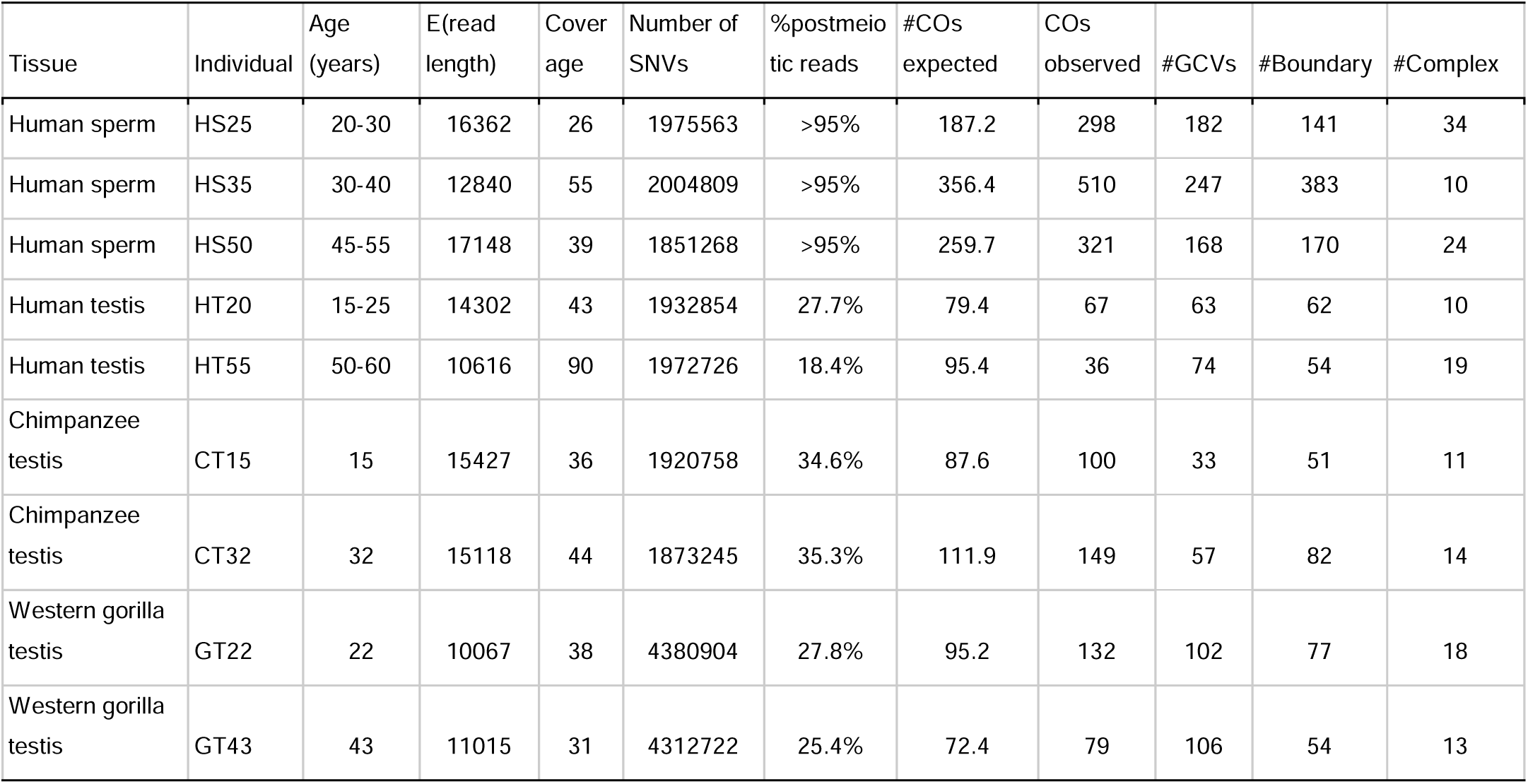
Identification of recombination events in sperm and testes samples. For human samples, age is anonymised to the nearest 5-year interval. The coverage is the mapped coverage of the haploid *de novo* assemblies of each sample before the classification of cell types. %post-meiotic cells are calculated from the classification of reads by CpG methylation patterns (see methods). From these, the expected number of COs are derived as coverage*(%post meiotic reads)*detectability*14. The last four columns are the number of the different recombination events identified in post-meiotic cells.

## Results

### Identification of recombination events from sperm and testis samples

We sequenced three human sperm samples purified by swim-up as well as testis specimens obtained from two individuals of humans, chimpanzees and western gorillas to 26-90X genomic coverage using Pacbio HiFi sequencing (Methods, Table 1). For each sample, we first constructed a high-quality *de novo* genome assembly (see Methods) covering more than 95% of the genome with N50 contig sizes of 50-93 Mb across samples (see Supplementary Table 1). We then mapped all long reads back to this assembly, identified all high-confidence single nucleotide variants (SNVs) and assigned variants to haplotypes, i.e. inferring the full phasing of all variants. Since reads typically have 10 SNVs, haplotype switch errors are exceedingly rare (see Methods). On this backbone, we interrogated all mapped reads for the occurrence of “variant shifts” between the haplotypes as a result of recombination events. After extensive filtering, we classified detected events into four types: COs, simple GCVs, complex GCVs and boundary cases (Supplementary Figure 1, Figure 1A). The short arms of the acrocentric chromosomes and segmental duplications have previously been shown to often engage in complex interallelic genetic exchanges^14^. We identified for the human sperm samples 17 (HS25), 29 (HS35) and 19 (HS50) events in acrocentric chromosomes and 83 (HS25), 105 (HS35) and 65 (HS55) in segmental duplications. However, for the present study, we have masked these regions for further analysis (Supplementary Table 5).

The testis tissue comprises a mixture of somatic cells (e.g. blood, peritubular, Leydig and Sertoli cells) and germline cells (diploid spermatogonia, tetraploid primary spermatocytes and diploid secondary spermatocytes, and haploid spermatids and spermatozoa). Here, we are only interested in postmeiotic events, and therefore, we need to identify the cellular origin of each sequencing read. It is possible to determine the methylation status on the majority of the 100- 200 CpG sites on an average PacBio HiFi read. We developed a classification method that, from the observed methylation pattern on a single PacBio HiFi read, estimates the probability of a read being of somatic or germline origin (see Methods). The method is trained on published methylation data from different cell types of human spermatogenesis^15^ and somatic proxies (blood and neurons). We can reliably classify more than 80% of the HiFi reads to somatic versus germline and even specific spermatogenic cell types (Supplementary Table 2). Based on this classification, we then restricted the analysis of putative recombination events to the germline. This is particularly important for GCV events where only a single SNV change between haplotypes occurs, as these can also be mimicked by recurrent mutations in somatic cells with higher mutation rates than germline cells^16,17^. Furthermore, it removes potential GCVs in cells that are not part of the germline.

We manually curated, via visual inspection, the remaining recombination events of all types using IGV with a high (>95%) interobserver concordance in the identification of false positives (see Methods). The number of likely false positives removed by this process varied between 7% and 77% (Supplementary Table 6). The curated set of events for each sample is shown in Table 1. We next estimated whether the number of COs identified agreed with prior expectations. We can estimate how many COs we expect to identify in each sample from the mapped sequencing coverage, the detectability of a CO given that it occurred on the read, and the estimated proportion of postmeiotic cells for the testis samples, as explained above. For COs, we have prior expectations of approximately 14 COs per haploid cell since the male genomic map length is around 28 Morgans^18^. For detectability, we estimated by simulation on each sample the probability that a random CO on a read will cause an SNV pattern as the one we use to identify COs (Figure 1A). The power of detection for CO events varies with the SNV density and read length and thus by sample. We estimate that across samples, between 30% and 52% of the COs should be detectable (Supplementary Figure 2). Multiplying these estimates and the estimated fraction of postmeiotic cells for the testes samples, we infer a number of expected COs for both sperm and testes samples (Table 1) that, given the many uncertainties, are close to the number of COs detected (10-30% differences except for the testis specimen from the elderly man, HT55). Part of this deviation could also be due to individual differences in the genetic map length. In HT55, we speculate that spermatogenesis is less efficient than estimated, i.e. a smaller proportion of the germ cells than expected are postmeiotic.

Likewise, the boundary cases, where an SNV at the end of a read changes haplotype, are expected to be a mixture of COs and GCVs (Supplementary Table 5) in proportion to their estimated frequency. We can estimate the expected number of these that are COs and GCV, respectively, and the sum of these matches well with the observed boundary cases (Supplementary Table 3). Together, these tests suggest that the detected sets of curated COs and GCVs are reliable.

### Genomic distributions of crossovers and gene conversions

Next, we investigated the spatial distribution of CO and GCV events across the genome (Figure 1B, sample HS35. The remaining samples are shown in Supplementary Figure 6). COs, in particular, are clustered at the telomeres, whereas GCVs are more uniformly distributed. We tested for a difference in clustering against a random distribution and found this to be highly significant (P<1e-05) in all samples except in the GT43 gorilla sample (P=0.05) and in the GT22 gorilla sample and HT20 human testis sample (not significant), which had few events (Supplementary Figure 6). We also investigated the density of COs and GCVs across each chromosome after all samples were pooled. Chromosome 2 was omitted from this analysis because of karyotype differences in humans versus great apes. Whereas the density of COs increases with decreasing chromosome size (Supplementary Figure 3A, P<0.001, linear regression of density of COs with chromosome size), the density of GCVs is not correlated with chromosome size (Supplementary Figure 3B, P=0.10). This results in an increasing CO/GCV with decreasing chromosome size (Supplementary Figure 3C, P=0.033). Thus, the larger recombination rate per base pairs on small chromosomes is due to a higher proportion of the DSBs resolved as COs.

Figure 1C shows that the observed number of CO events per Mb decreases rapidly away from the telomere for each of the four types of samples (human sperm, human testis, chimp testis and gorilla testis pooled). This effect is much weaker for inferred GCV events.

Since a male pedigree recombination map also reveals higher recombination rates near telomeres, we related the positions of COs to the deCode recombination map (smoothed at a 100 kb scale). Figure 2A,B shows that the positions of the COs we identify are indeed highly enriched in regions with higher estimated recombination both for human sperm and human testes samples. GCVs are also enriched but to a much smaller extent, as expected, since COs are the basis for the recombination map.

**Figure 2:**
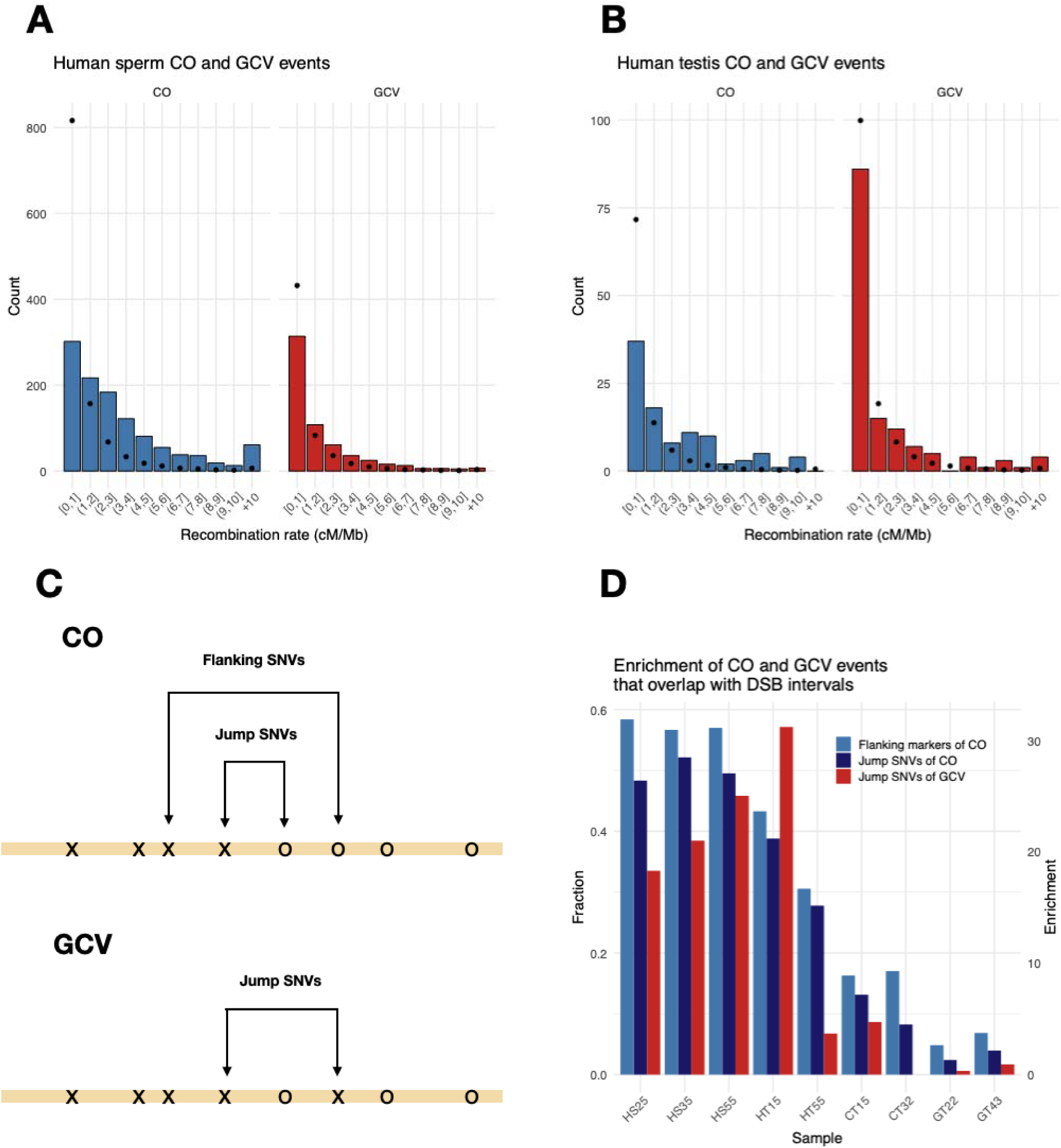
Double strand breaks and inferred recombination events. **A.B.** The recombination rate (deCode map at 100 kb scale) for observed CO and GCV events pooled by human sperm cells samples and human testes, respectively. Histograms indicate observed counts, while dots denote expectations under a uniform distribution along the genome. B. Schematic illustration of flanking SNVs and jump SNVs for COs and GCVs. C. The fraction of crossovers and gene conversions in hotspots of DSB and the enrichment of crossovers and gene conversions in hotspots of DSB ^1^.

Since COs and GCVs occur after programmed DNA DSBs, our results suggest that a non-random set of telomere proximal DSBs is resolved as COs. To test this more directly, we lifted a map of DSB hotspots from spermatocytes^1^ to the T2T-CHM13v2.0 genome and studied the overlaps with the positions of our detected COs and GCVs. We separately recorded how many DSBs overlapped with the region between the jump SNVs (the polymorphisms flanking a CO event or a tract of a GCV event) and also a larger region, including flanking SNVs of the CO since we, from the biased GCV results discussed below, suspected that some COs could be initiated outside of the jump SNVs (see Figure 2C). We find that for human samples, 40-60% of COs and GCVs overlap with a DSB hotspot, leading to an enrichment of around 30-fold compared to a random overlap. This enrichment is higher when including the flanking SNVs, supporting that some COs are initiated outside of the region marked by the jump SNVs but one SNV is then gene converted in the process (Figure 2D). Again, the positions of GCV events called in the testis sample of the elderly man, HT55, deviate from this pattern.

We do not see a similar strong enrichment between the human DSB hotspots and CO and GCV events for chimpanzees and western gorillas, suggesting that the DSB hotspots are not conserved between great ape species. This is expected since DSB placements are directed by PRDM9 motifs, which differ between the three species^19,20^. Interestingly, the events called in the more closely human-related chimpanzee samples show some enrichment (CT15 and CT32, Figure 2D), suggesting a greater overlap in DSBs between humans and chimpanzees.

### Gene conversion tract lengths and rates

Since NCO tracts are expected to be short and the density of SNVs is low, most NCO events do not move SNVs, and most of the inferred GCVs only move one polymorphic site (83-92% across samples, see Supplementary Table 3). Thus, instead of averaging the length of events detected, we used all data in each sample to infer the lengths of GCVs from a probabilistic model. We derived a likelihood model that infers an average tract length for GCVs from the number of GCVs moving successive (1, 2, 3…) polymorphic sites and the observed distances between heterozygous sites in the genome of the individual investigated. The model assumes that the NCO tract length is geometrically distributed (see ^13,21,22^ and Charmouh et al., BIORXIV/2024/601865, for model details). In all samples, the estimated mean tract lengths are 40-100 bp (Figure 3A). A likelihood ratio test could not reject the hypothesis of equal mean tract lengths across the samples (Supplementary Figure 5). The short estimated tract lengths imply that across the genome, only 2-5% of NCOs move at least one SNV and hence are detectable as GCV events (Figure 3B, Supplementary Table 3). We combined our estimates of mean tract length and rates of detection to estimate the overall probability that a bp is part of an NCO event in meiosis (Figure 3C, Supplementary Table 4). Our estimates of 2-5 bp converted per 1 million bps align with previous estimates ^13,23,24^ and suggest variation among individuals.

**Figure 3.**
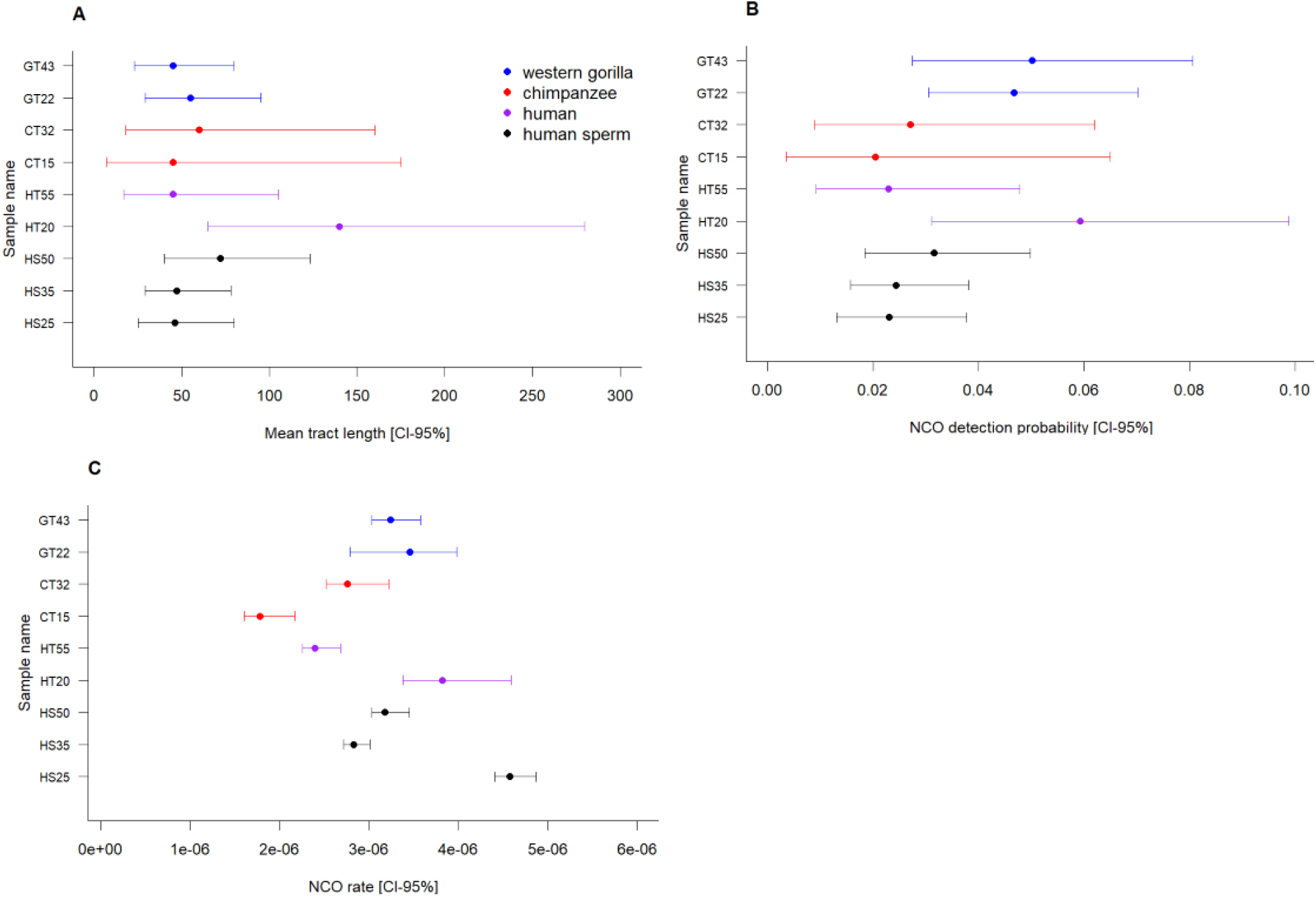
Gene conversion properties. **A.** Estimated mean tract lengths (error bars denote 95% confidence intervals) from the empirical SNV density and the numbers of SNVs moved in gene conversion events. **B.** The estimated fraction of NCO events that result in a gene conversion event, i.e. move one or more adjacent SNVs from one haplotype to the other. **C.** The estimated rate of gene conversion, i.e. the fraction of nucleotides involved in an NCO event in one generation

These results are based on simple GCV events alone. We also observe 10-34 complex events per sample that switch haplotypes several times (Table 1). These potentially represent more complex GCVs and/or COs with GCVs that are associated with longer NCO tracts than the simple GCVs discussed above.

### Biased gene conversion

Strand invasion associated with DSB repair will cause heteroduplex formation when the homologous chromosomes differ in the form of SNVs or indels. If such heteroduplexes contain both a strong (G,C) and a weak (A,T) allele, the repair is typically biased towards the strong base pairs. Such gBGC is a pervasive evolutionary force that shapes GC content in the genome and is the cause of a universal positive correlation between recombination rates and GC content in genomes ^7,9^. We examined the degree of GC bias in the different types of recombination events separately. First, for simple GCVs, we investigated the GC bias for events occurring in germline cells separately for single SNV and multiple SNV events since they have previously been shown to differ in GC bias in mice^21^. Overall, the human sperm data shows significant gBGC (P=0.00058, binomial test). For single SNV GCVs, we find an average gBGC of 54.6%+/- 2.1% (P=0.027, binomial test) (Figure 4A) and a stronger gBGC of 66.6%+/-4.2% (P=0.00034, binomial test) for multiple SNV GCVs (Figure 4B). This is in contrast to the results for mice, where multiple SNVs were found to be unbiased. For the testis samples, the results are more mixed with an overall gBGC observed only for multiple SNV events.

**Figure 4:**
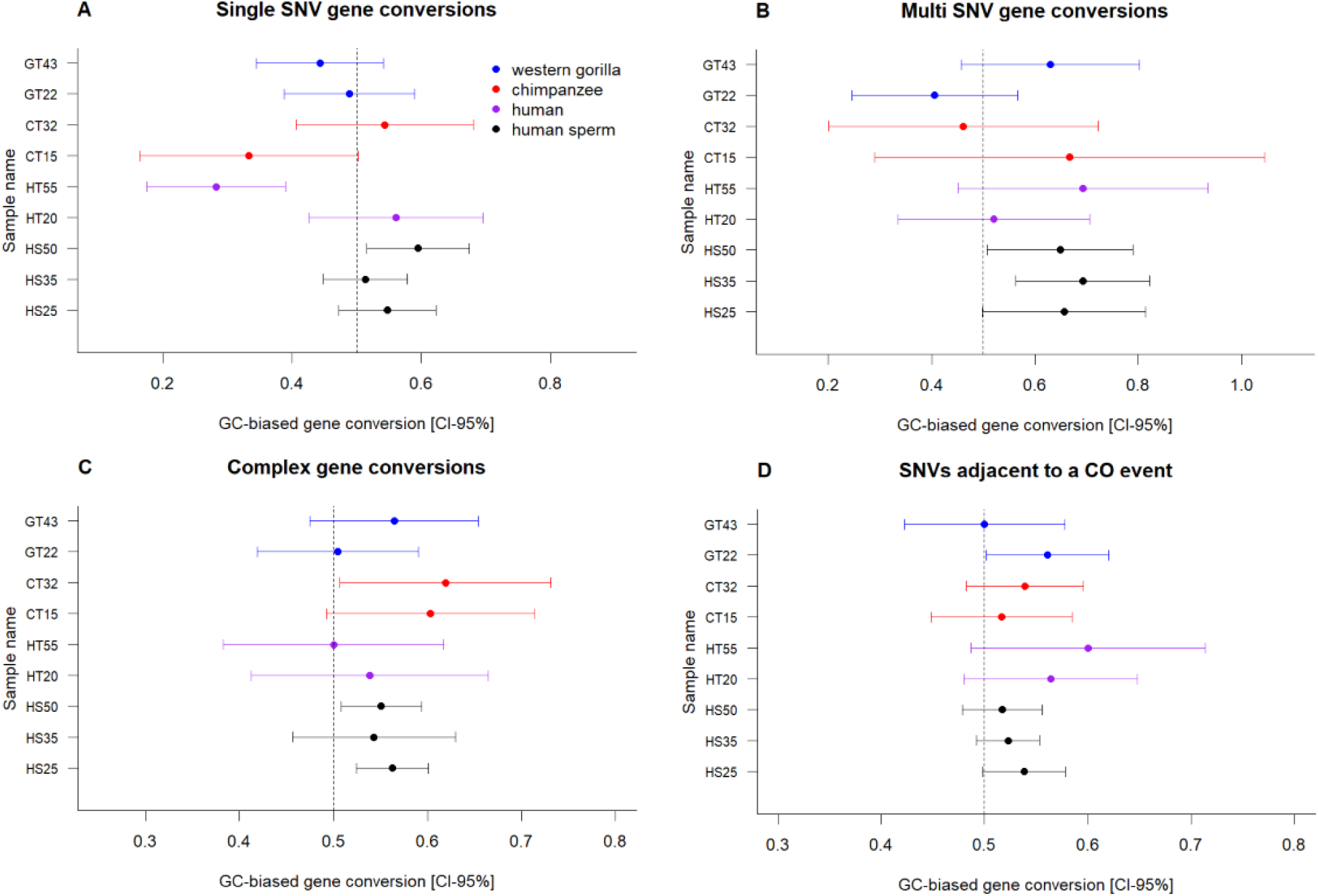
Estimates of the amount of gBGC with binomial 95% confidence intervals. **A.** Gene conversions where a single SNV is moved from one haplotype to the other. **B.** Gene conversions are where more than one subsequent SNV is moved. **C.** SNVs in a complex gene conversion event. **D.** SNVs adjacent to an inferred crossover point, the jump SNVs (see Figure 2B).

Next, we tested whether SNVs immediately flanking crossover events also showed signs of gBGC. We find a bias for all samples, with a significant bias for human sperm samples combined 52.5%+/-1.0% (P=0.014, binomial test) and for testis samples combined 54.1%+/- 1.5% (P=0.016, binomial test) (Figure 4C). This suggests that NCOs associated with COs are either strongly biased or that this type of NCO is much longer than the simple NCOs since the distance between SNVs is much longer than the estimated NCO tract length above (1.5 kb versus 40-100 bps). This agrees with previous estimates of NCOs associated with COs of about 500 bps ^21^. As a control, we also tested SNVs two positions away from the estimated CO point and found these not to be GC-biased (Supplementary Figure 5). These results suggest that some CO events indeed initiate outside of the interval marked by the SNVs that move from one haplotype to another. This also explains why we found a further enrichment of DSBs when including the region with the flanking SNVs of the jump SNVs (Figure 2C). Finally, we also investigated all SNVs participating in the complex GCV events for gBGC by just recording the percentage of weak to strong base pairs for all SNVs. Again, we find a consistent bias suggesting that these events, though relatively rare, will also contribute to GC content evolution (Figure 4D)

## Discussion

Our results confirm that in humans, DSBs induced by PRDM9 are responsible for most recombination events. A smaller fraction of these (5-10%) are resolved as COs, and this process is biased towards the telomeres and smaller chromosomes. This implies that GCV events are less correlated to pedigree-based recombination maps. In chimpanzees and Western gorillas, the overall telomeric enrichment of COs is also observed. However, at a finer scale, corresponding to locations of DSBs in humans (average size 2 kb), the overlap is very limited in agreement with both Western gorilla and chimpanzee genomes harbouring divergent PRDM9 motifs from those reported in humans ^25,26^.

Our estimates of NCO tract lengths from simple events are remarkably similar among species and tissue types, with no cases of significant differences. The estimated tract lengths associated with NCO events are short (averages between 40-100 bp), and they conform closely to a geometric distribution. In contrast, from gBGC patterns, we estimate that tracts of exchange associated with COs are much longer, at least on the order of 500-1000 bps. This may also explain the few cases of long complex GCV also associated with gBGC. We cannot determine the maximum length of these due to the finite read lengths, but they could be the rare fraction of long NCO events recently estimated by ^22^. Thus, the main effect of gBGC on GC content may not be from the resolution of simple NCO events but rather these longer tracts associated with COs or complex events. This would explain why GC content correlates very well with pedigree-based recombination rates, even though NCO events do not correlate very well with recombination rates.

We have shown that Pacbio HiFi reads allow precise mapping of COs and GCVs in post-meiotic cells of testis biopsies and in sperm samples. The ability to predict CpG methylation using HiFi kinetics could be leveraged to classify the reads coming from postmeiotic germline cells reliably. This is important since events mimicking both COs and GCV, in particular, are likely to occur in somatic cell types. We can exploit the striking patterns of methylation during spermatogenesis to identify with high confidence Pacbio HiFi reads that specifically originate from postmeiotic germline cells. This means recombination can now be studied from testis biopsies. This opens the possibility of studying recombination in a much broader range of individuals and or species where extensive trios or sperm samples are not currently available.

## Methods

### Samples

DNA was isolated from human testis tissue obtained from two anonymous men (autopsies) who consented to donate tissue for research purposes post-mortem. Ongoing spermatogenesis with post-meiotic round spermatids in most tubules was verified by histological staining. The study also included human sperm samples that were obtained from three anonymous sperm donors with good semen quality. The sperm samples were obtained in 2018 after consent and an ethical permit granted by the regional committee (H-17012149). As human tissue and sperm donors were anonymous the exact age was unknown, but an approximate estimate given. Finally, the study included testis tissue from four great apes, two chimpanzees of the West African subspecies (*Pan troglodytes verus*; CT15 is European Association of Zoos and Aquaria (EAZA) studbook number 13342 and CT32 is EAZA studbook number 12295 and two Western gorillas of the Western lowland subspecies (*Gorilla gorilla gorilla*); GT22 is EAZA studbook number 1435 and GT43 is EAZA studbook number 492. The samples from great apes housed in zoos within the EU were collected post-mortem after death by natural causes. Samples are named according to species (H for human, G for gorillas, C for chimpanzees), tissue (T for testis and S for sperm) as well as the age (estimated for human samples).

### DNA extraction

DNA was isolated from testis tissue samples using the MagAttract® HMW DNA kit (Qiagen, Hilden, Germany) following the manufacturer’s instructions. The three sperm samples were subjected to gradient purification using PureSperm 50 (Nidacon, Gothenburg, Sweden) to avoid somatic cell contamination and enrich for mature sperm. The fraction containing mature sperm was then incubated with RLT buffer (Qiagen) and stainless steel beads on a shaker for 10 minutes to allow better access to the tightly packed DNA. The DNA was subsequently isolated using the automated Maxwell 16 system (SEV AS1010, Promaga, Madison, WI, USA).

The quantity and quality of the isolated DNA were evaluated by Nanodrop (ThermoFisher Scientific, Waltham, MA, USA), Qubit dsDNA HS or BR Assay Kits (ThermoFisher), and gel electrophoresis before library construction. All samples revealed DNA of high-molecular weight.

### PacBio HiFi sequencing

The DNA was subjected to PacBio HiFi sequencing at the Norwegian Sequencing Centre (www.sequencing.uio.no), a national technology platform hosted by the University of Oslo and supported by the “Functional Genomics” and “Infrastructure” programs of the Research Council of Norway and the Southeastern Regional Health Authorities.

In short, the sequencing libraries were prepared using Pacific Biosciences protocol for HiFi library prep using SMRTbell® ExpressTemplate Prep Kit version 2.0 (Pacific Biosciences, Menlo Park, CA, USA). DNA was fragmented into 15-20 kb fragments using Megaruptor 3 (Diagenode, Denville, NJ, USA), and the final library was size-selected using BluePippin (Sage Science) with a 10 kb cut-off.

The final libraries were sequenced on the Sequel II instrument using 8M SMRT cells with a 30h movie time and the Sequel II Binding kit 2.2 and Sequencing chemistry v2.0 (Pacific Biosciences).

Four samples (HT55, GT43, GT22, HS35) were, in addition to sequencing on the Sequel II instrument, also sequenced on the newer Revio instrument using SMRTbell® ExpressTemplate Prep Kit version 3.0, 25M SMRT cells, the Revio Polymerase kit and a 24h movie time. A single sample (HS25) was only sequenced on the Revio instrument using the same kits and parameters.

CCS sequences were generated using CCS pipeline (SMRT Link v10.2.0.133434 for Sequell II and v. 12.0.0.183503 for Revio reads) and reads with at least 99% accuracy retained as HiFi reads.

### De novo assemblies

The *de novo* assemblies were built with *hifiasm* with its default parameters. The summary statistics of the assemblies are shown in Supplementary Table 1.

### Mapping

Following the construction of the de novo assemblies, we map the reads from each individual against its de novo assembly using *pbmm2* with parameters -c 99 and -l 2740.

### SNV calling

The SNVs were called using an in-house Python script based on pysam. The program iterates through a BAM file and stops at a position where exactly two different nucleotides are present. Furthermore, the coverage in this position has to be greater than the 5th percentile and less than the 99.7th percentile of the coverage distribution. The program also requires that no more than 10 % of the reads can contain an indel at the given position and that the variant occurs in at least three reads.

Since PacBio HiFi reads accuracy is lower in repetitive regions, we filtered candidate SNVs that fall within local repetitive regions. Specifically, if the number of unique 4-mers in a 32-bp region is less than 19, we skip the position. We also skip the position if the candidate SNV occurs at the boundary of two homopolymers. For instance, if the sequence context is AAAA(T/A)TTTT, then it is difficult to assess whether the SNV is real or due to a sequencing artefact.

The last criterion for a position being an SNV is that it cannot occur within 15 bp of an indel column. We define an indel column as a position where more than 10 % of the reads contain an indel.

For all reads at an SNV position, we assign a haplotype to each read. If the read contains the same base as the de novo assembly, the read is assigned haplotype ‘X’, whereas the read is assigned haplotype ‘O’, if it contains the alternative nucleotide. In rare cases, some reads contain an indel at the SNV position, and these reads are assigned haplotype ’n’. If a read is assigned ‘O’, but the base quality is less than QV35, or a small indel is within 10 bp in each direction, or a mismatch is within 10 bp in each direction, or a big indel (>10 bp) is within 250 bp in each direction, then the read is assigned haplotype ‘o’. The distinction between ‘O’ and ‘o’ makes it possible to assess the quality of the haplotype assignment for all reads at all SNV positions. An identical assessment is performed for haplotype ‘X’. Following the assignment of haplotypes, the program continues until the last position in each contig.

### Recombination calling

Candidate recombination events are called when a read contains high-confidence variants from both haplotypes (X and O). Subsequently, those events are divided into four basic types (see also Supplementary Figure 1):

1. CO events are defined as reads that display a single jump of haplotype and where at least two SNVs from each haplotype are present (e.g., ‘XXxOO’).
2. NCO events are defined as an SNV string that contains exactly two haplotype jumps (e.g. ’OoXOO’).
3. The boundary events are defined as an SNV string with a single jump of haplotype (like the COs), but the jump occurs between the first or last two SNVs (e.g., OOooX).
4. The complex events are defined as an SNV string with more than two haplotype jumps. To minimise the number of false positives, we exclude events that occur closer than 5000 bp to another event.

The overall pipeline is sketched in Supplementary Figure 1, however the final step in filtering the reads containing recombination events is specific to the human samples, where we exclude any recombination calls that land on the acrocentric short arms, and any calls that overlap with segmental duplications.

To separate the events found in segmental duplications, we intersect the map of segmental duplications with the CHM13 coordinates of the called events ^27^.

SD and acrocentric annotations: https://zenodo.org/records/7671779

Furthermore, for testis samples, we only record the recombination events observed in reads that are assigned as germline reads in Figure 1A. For the classification of reads, see the section “Methylation-based classification of cell types” below.

### Manual curation

Following the classification of germline read recombination events, all reads were manually curated using a script rendering relevant IGV sections. Manual curation was done by PSP for all events and by APC, SB, and MHS for a subset of the events. From this, we calculated the interobserver concordance to be >0.95, with 5-20% of events deemed as false positives and removed from further consideration. The events scored as false positives were mainly due to repetitive regions of the genome, long distances between SNVs, or collapsed regions, where more than haplotypes are present.

### Inference of distribution of NCO tract length from candidate read counts

We estimated NCO tract lengths using the framework of (Charmouh et al 2024 BIORXIV/2024/601865). Briefly, we infer, using maximum likelihood, the mean of the best fitting geometric distribution of NCO tract length from the observed counts of reads that exhibit a footprint of NCO and by tallying how many reads convert/move 1, 2, 3 etc. SNVs in the read from an O to an X haplotype. This approach assumes that all NCO events are independent and that each NCO event induces a tract of physical length L, where L is modelled as a stochastic (random) variable geometrically distributed with expected mean 1/s. The data for each sample is summarised as the counts of reads n_i containing apparent GCV events that have moved in SNVs.

Using simulation, the method allows us to obtain the expected proportions of reads harbouring 1,2,3, etc. converted SNVs given the genome-wide SNV distribution of each sample as a function of some mean NCO tract length. By doing so, we account for differences in detectability induced by the fact that the SNV density underlying the two haplotypes varies hugely from region to region (Median distance: 347 bp, mean inter SNV distance: 1173.07 bp, IQR: 812, see Supplementary Figure 7).

### Testing for differences in NCO tract length between samples

To test for differences in tract length between samples A and B, say, we employ a likelihood ratio test comparing the relative fit of two models to the data. Under Model0 (M_0_), we assume that reads from both samples are independent observations of NCO where the induced conversion tract length is drawn from a single geometric distribution with mean length L_0_ = 1/S_0_. Under Model1 (M_1_), we allow the induced conversion tract lengths to follow different geometric distributions with mean 1/S_A_ and 1/S_B_, respectively. The likelihood of the data **D** (counts in sample A and sample B) are maximised under both M_0_ and M_1_, and we use G_obs_ = 2 log (L_1_(**D**)/(L_0_(**D**)) to assess statistical significance. Under the null hypothesis that both samples follow an identical tract length, we have that G_obs_ is approximately LJ^2^ distributed with 1 degree of freedom.

Using this framework, we can test for pairwise differences (e.g., between sample A and B) or extend the test to globally test for differences between all K samples. In that case, M_1_ allows K geometric distributions with different means for each sample, and the test statistics G_obs_ is approximately LJ^2^ distributed with K-1 degrees of freedom if M_0_ is correct.

Given that the likelihood functions rely on the multinomial distribution of counts, and provided that the total number of counts is larger than 5, the LJ^2^ approximation to the distribution of G_obs_ is expected to be very accurate (e.g.^28^, Chapter 8).

### Methylation-based classification of cell types

To develop a binomial-based method for reads classification into germline or non-germline, we required binomial distribution parameters for all CpG sites of the reference genome in germline and non-germline cell types separately.

Hence, we obtained the methylated and non-methylated counts at individual CpG sites of the reference genome (hg38) in germline cells using genome-wide methylation data originating from NEBNext Enzymatic Methyl-seq (EM-seq; New England Biolabs, Ipswich, MA, USA) of flow-sorted spermatogenic cell types representing 4 different stages of spermatogenesis. The dataset contained 4 sorted spermatogenic cell types obtained from 3 different men with ongoing spermatogenesis and represent undifferentiated spermatogonia (2C, UTF1+/DMRT1-), differentiating spermatogonia (2C; UTF1-/DMRT1+), primary spermatocytes (4C) and spermatids/spermatozoa (1C)^15^. Finally, WGBS data from ejaculated and isolated sperm from 5 men were also included^29^

Similarly, to obtain methylation counts for non-germline or somatic cells, we used whole-genome bisulfite sequencing (WGBS) samples from cell-sorted blood samples and brain biopsies from multiple individuals^29^. For blood and brain tissue, the different cell types and samples were grouped to form one high-coverage estimate of the methylation levels for each CpG site in the blood (based on 10 B-cells, 36 T-cells, 6 Monocytes, 4 Natural Killers, and 4 Granulocytes files) and brain (based on 20 neuron files).

We thus consider 7 different types of cells: 5 different types of germline cells corresponding to mature sperm plus the four spermatogenic cell stages of spermatogenesis and two types of somatic cells (blood and brain).

All sequencing reads of the training data were aligned to hg38, and reference genome CpG sites with a coverage smaller than six (12.3% of all CpG sites of the genome) were discarded. For each CpG site, *i*, we counted the number of methylated (n_m,i,c_) and unmethylated (n_u,i,c_) reads in each of the seven cell types, c The HiFi reads we wanted to classify were also aligned to hg38. For each CpG site, i, in a read, r, a methylation status, *p_r,i_*, ranging from 0 (unmethylated) to 1 (methylated) had been calculated using jasmine and we use this as our raw input to build our classifier for each read. https://github.com/PacificBiosciences/jasmine.

Our analysis excluded CpG sites on a HiFi read with a high degree of uncertainty about methylation status (*p_r,i_* between 0.3 and 0.7) and CpG sites that were not present in the training data CpG sites.

We can then calculate the log-likelihood of observing this *p_r,i_*at site, i, given that read, *r*, comes from cell type, *c*, assuming that the methylation level in the different cell types follows a Beta distribution. For each type of cell, we use a beta distribution empirically motivated by our training data, e.g. the observed counts of methylated (n_m,i,c_) and unmethylated (n_u,i,c_) reads in each type of tissue c. :

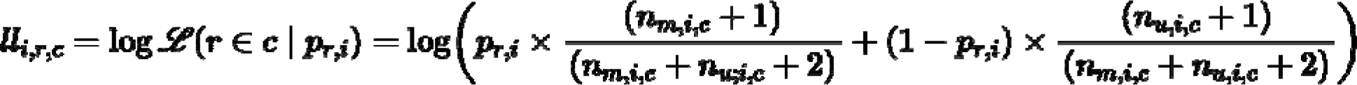

To estimate the log-likelihood that read, *r*, comes from cell type, *c*, using the methylation status 671 of all informative CpG sites on the read. We assume independence between the methylation 672 levels of each CpG site and, accordingly, sum the site-specific log-likelihoods. If the read 673 contains *L* CpG sites starting with number x this becomes:

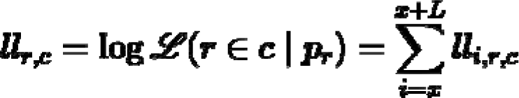

We then classify a read as germline if the cell type with the highest log-likelihood is one of the five germline cell types.

## Acknowledgements

The authors are grateful to Bonnie Colville-Ebeling for helping with the collection of anonymous human testis tissue and Brian Vendelbo Hansen for helping with the DNA purification. We are also grateful to Veterinarian Imke Lüders, Münster Zoo, Germany for helping collect great ape samples and Senior Engineer Ave Tooming-Klunderud from the Norwegian Sequencing Centre for the excellent help with the PacBio HiFi sequencing. The computational work has taken advantage of the genomeDK supercomputing infrastructure. The work was funded by the Novo Nordisk Foundation (grant# NNF21OC0069105). The authors have benefited from networking grants of COST Action CA20119 (ANDRONET) supported by the European Cooperation in Science and Technology (www.cost.eu).

## Author contributions

MHS, TB, SB and KA conceived the study. CH, KA and SBW collected the samples. SBW and KA generated the data. PSP built the de novo assembly, called recombination events, analysed the spatial distribution of recombination events and headed the manual curation of reads with input from MHS, TB, APC and SB. APC developed and applied inference methods for estimating gene conversion and gBGC with input from PSP, MHS, TB, SB and AH. VKS and SB developed and applied the read classifier. MHS and PSP wrote the first draft of the manuscript with substantial input from APC, TB, AH, KA, SB, VKS, SBW, MP and SL.

## Data availability

All scripts used for analysis are available at [https://github.com/PeterSoerud/recombination_calling]. According to Danish legislation, we are not allowed to deposit human sequencing data from anonymous human donors. For chimpanzees and gorillas, HiFi reads in fastq format have been deposited in the ENA archive, accession number PRJEB77177

---

**Supplementary Figure 1.**
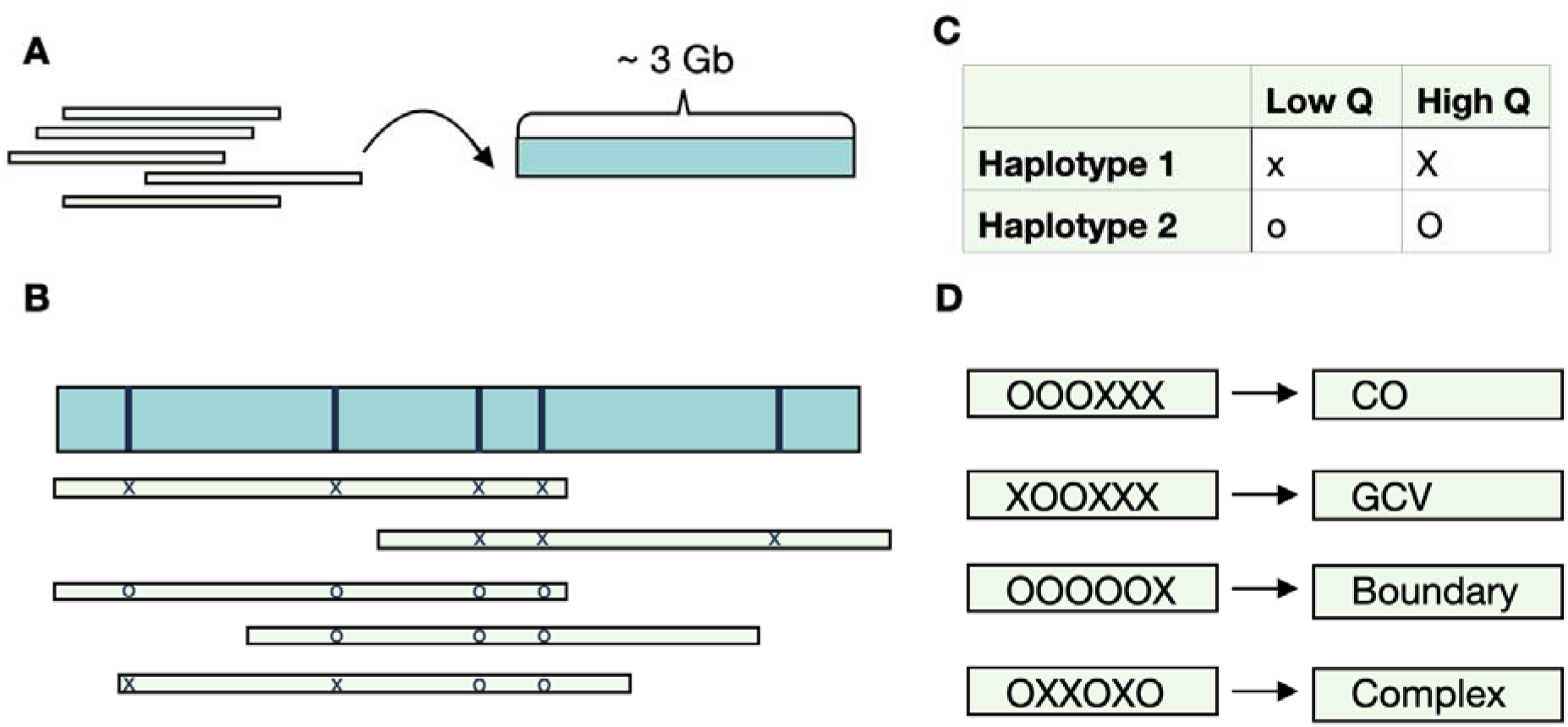
Schematic pipeline for the identification of recombination events. **A.** From the raw HiFi PacBio reads, a high-quality *de novo* assembly with N50 contig size >50 Mb was constructed for each sample. B. Mapping each individual read back to the *de novo* assembly, high-confidence heterozygous positions (vertical bars) were identified along the assembly and fully phased as Haplotype 1 and Haplotype 2 (not shown here). The sequence of single nucleotide variants at each heterozygous positions (marked as X/x’s and O/o’s) were used to assign unambiguously each individual read to a parental haplotype, conversely, to identify candidate reads harbouring shifts between the two haplotypes that indicate a potential recombination event (such as the bottom read with the sequence XXOO). C. Each mapped read was interrogated for the occurrence of shifts between the two parental haplotypes using a set of stringent quality thresholds on base quality, mapping quality, presence of indels, and soft- clipping. D. The candidate reads were then classified into four different types of recombination events based on the type of shifts between haplotypes.

**Supplementary Figure 2:**
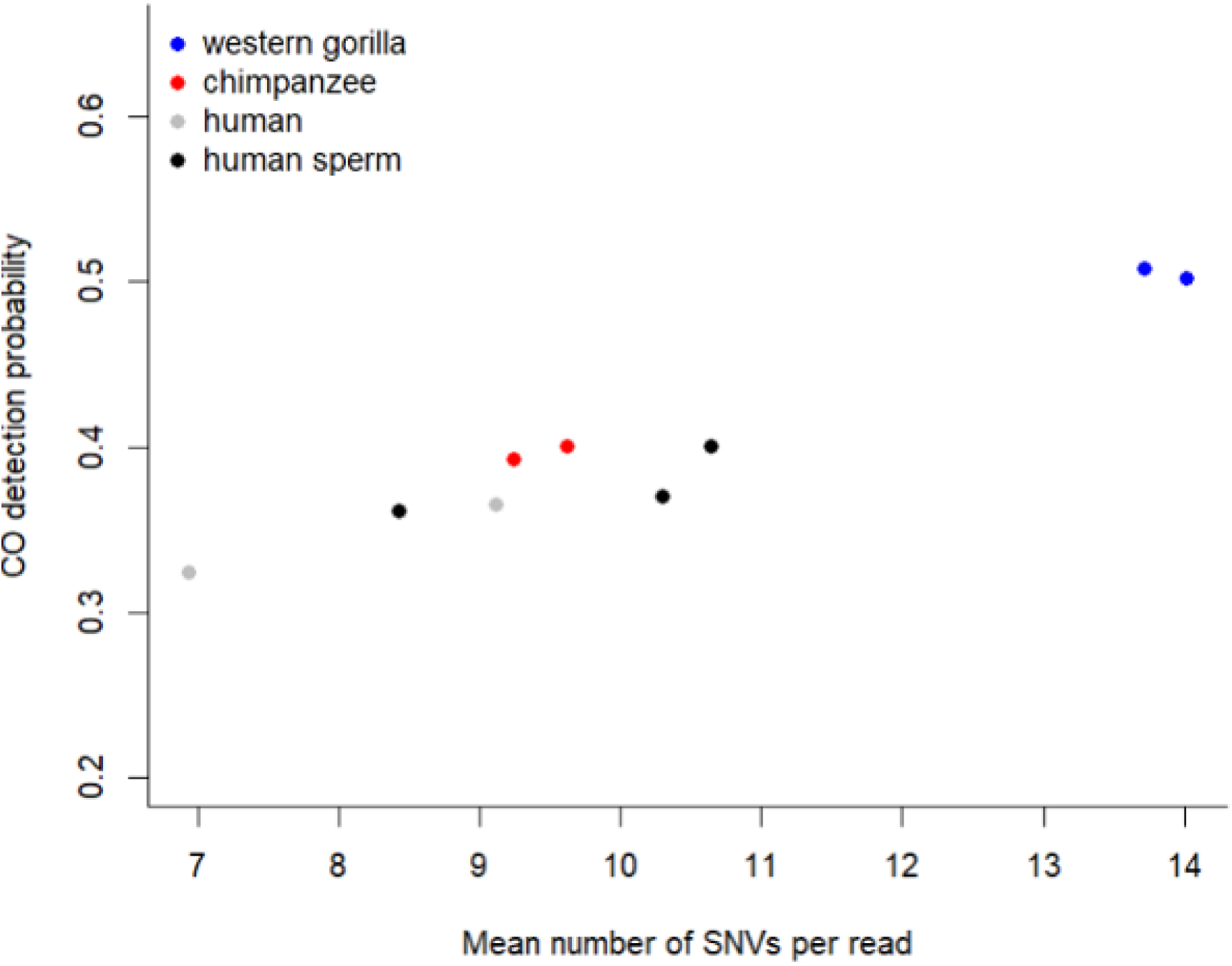
Crossover (CO) detectability as a function of the mean number of SNVs per read. Since calling a CO event requires 2 flanking SNVs on either side of the event, the product of mean read length and SNVs density (i.e. mean number of SNVs per read) explains most of the variance in CO detectability among samples.

**Supplementary Figure 3:**
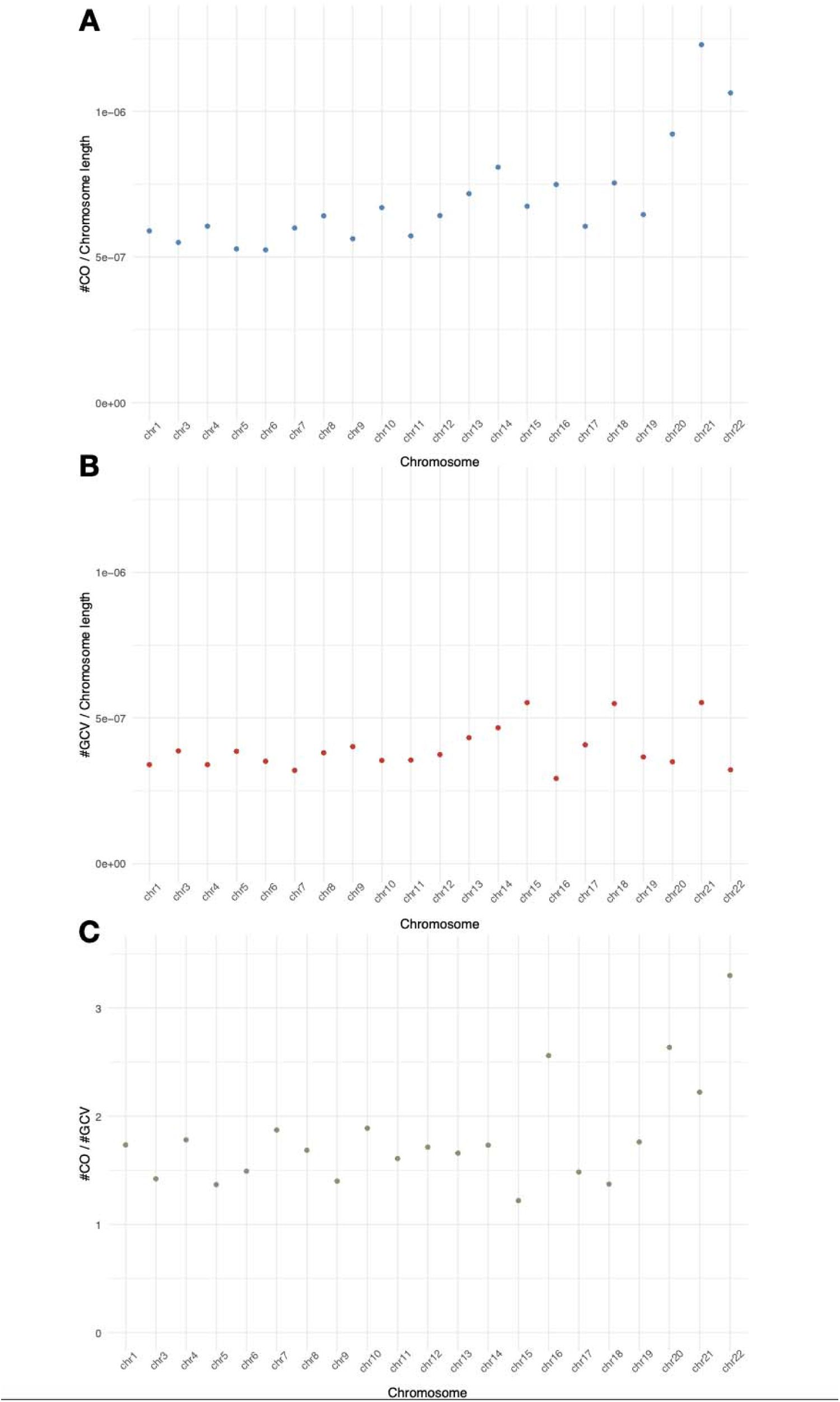
Chromosomal distribution of recombinations. A. The probability of a crossover per bp per chromosome (chr) when combining all events across samples (chr 2 omitted). B. The probability of a gene conversion per base pair per chromosome pooling all events across samples (chr 2 omitted). C. The ratio of crossover to gene conversion per chromosome shows an increase with decreasing chromosome size (P=0.03, linear regression of ratio with chromosome size).

**Supplementary Figure 4:**
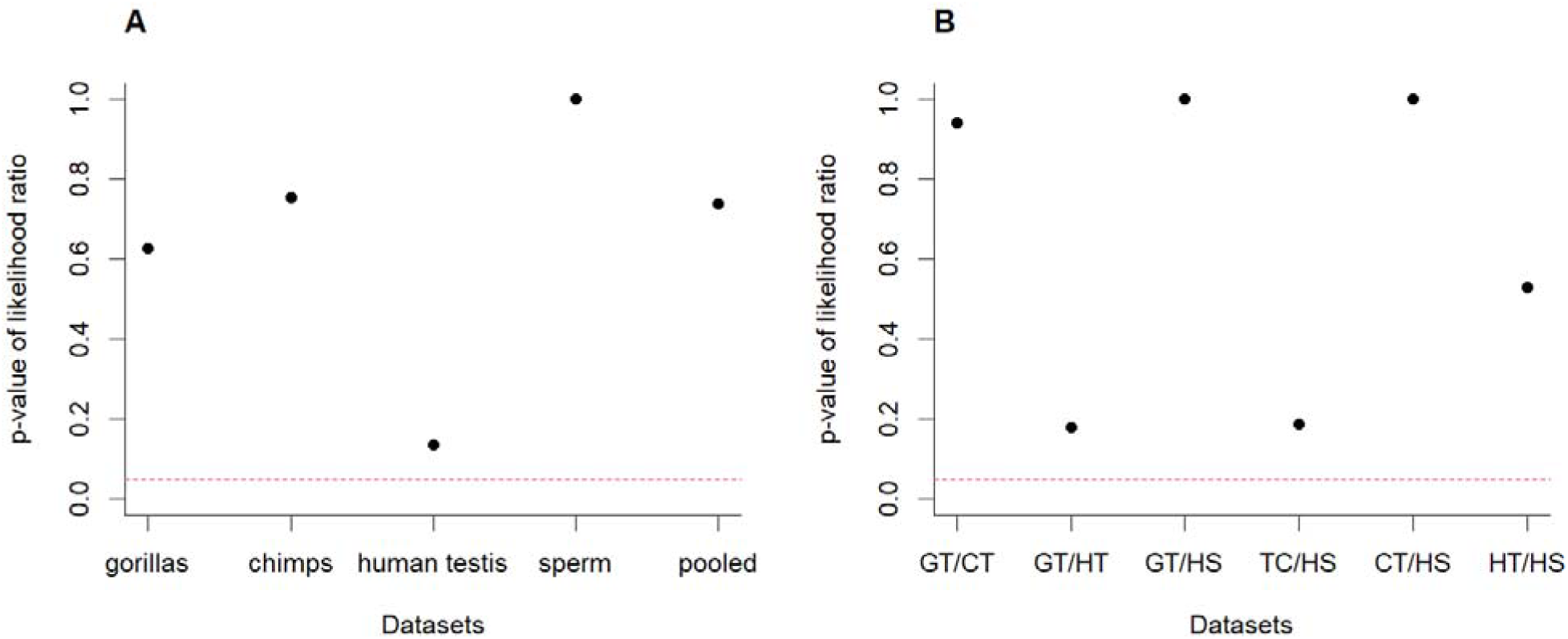
P-values for the likelihood ratio test of whether a single mean tract length fits multiple datasets significantly better than different mean tract lengths for each dataset (see methods for details). The dashed red line shows a significance level of 0.05. Despite the p- value not being adjusted for multiple testing, there are no significant differences in the likelihood of models explaining (A) a specific tissue type with a single mean tract length or all tissues with a single mean tract length (pooled) versus a specific mean tract length for each sample. Similarly, there are no significant differences in likelihood between models describing two tissue types with a single mean tract length versus models assigning a mean tract length to each tissue type (B). This suggests that the mean tract length across the 3 different species we examined is evolutionarily conserved.

**Supplementary Figure 5:**
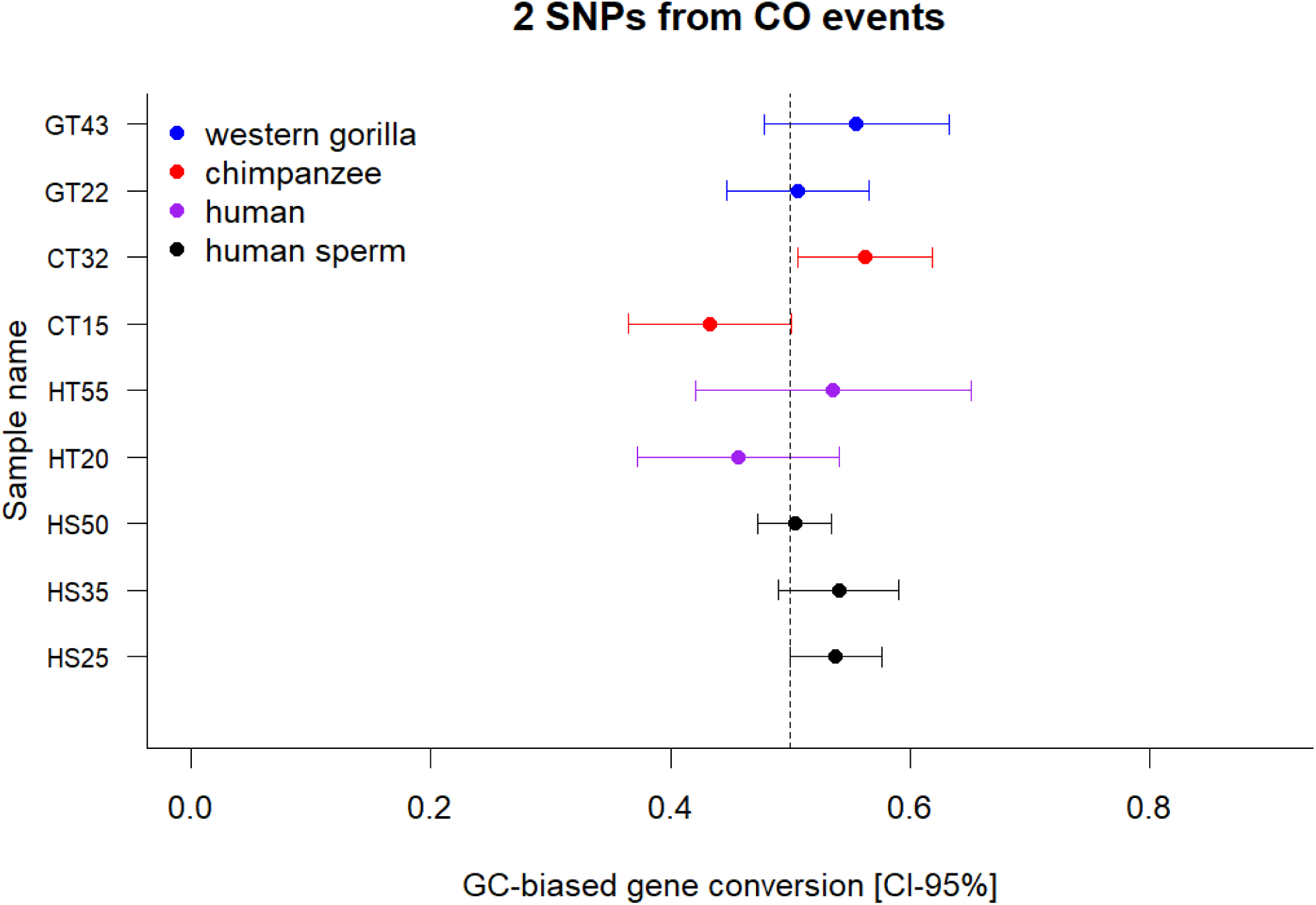
No indication of GC-biased gene conversion for the flanking SNVs of a recombination, i.e. positioned next to the jump SNVs (Figure 2B).

**Supplementary Figure 6:**
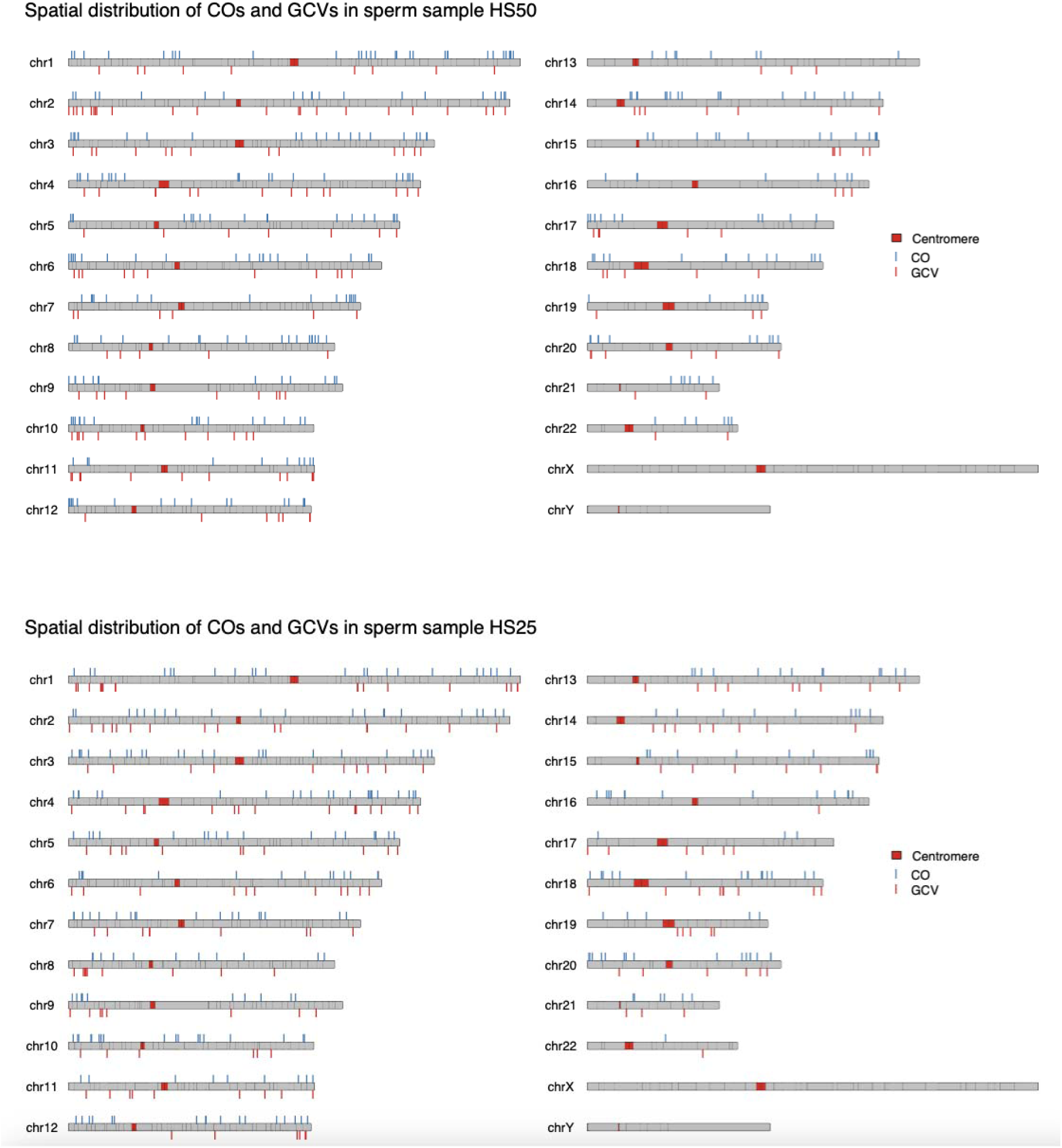
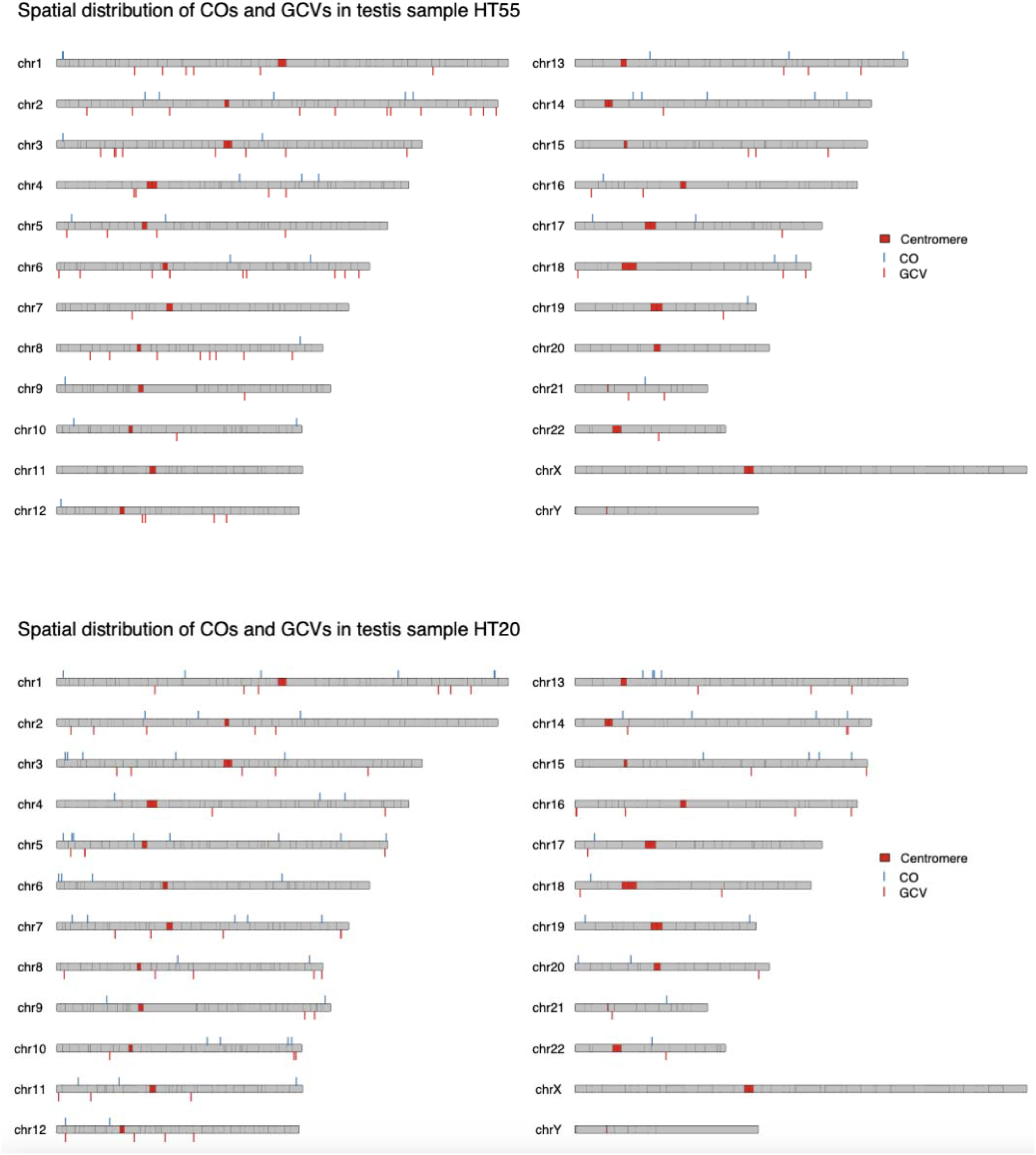
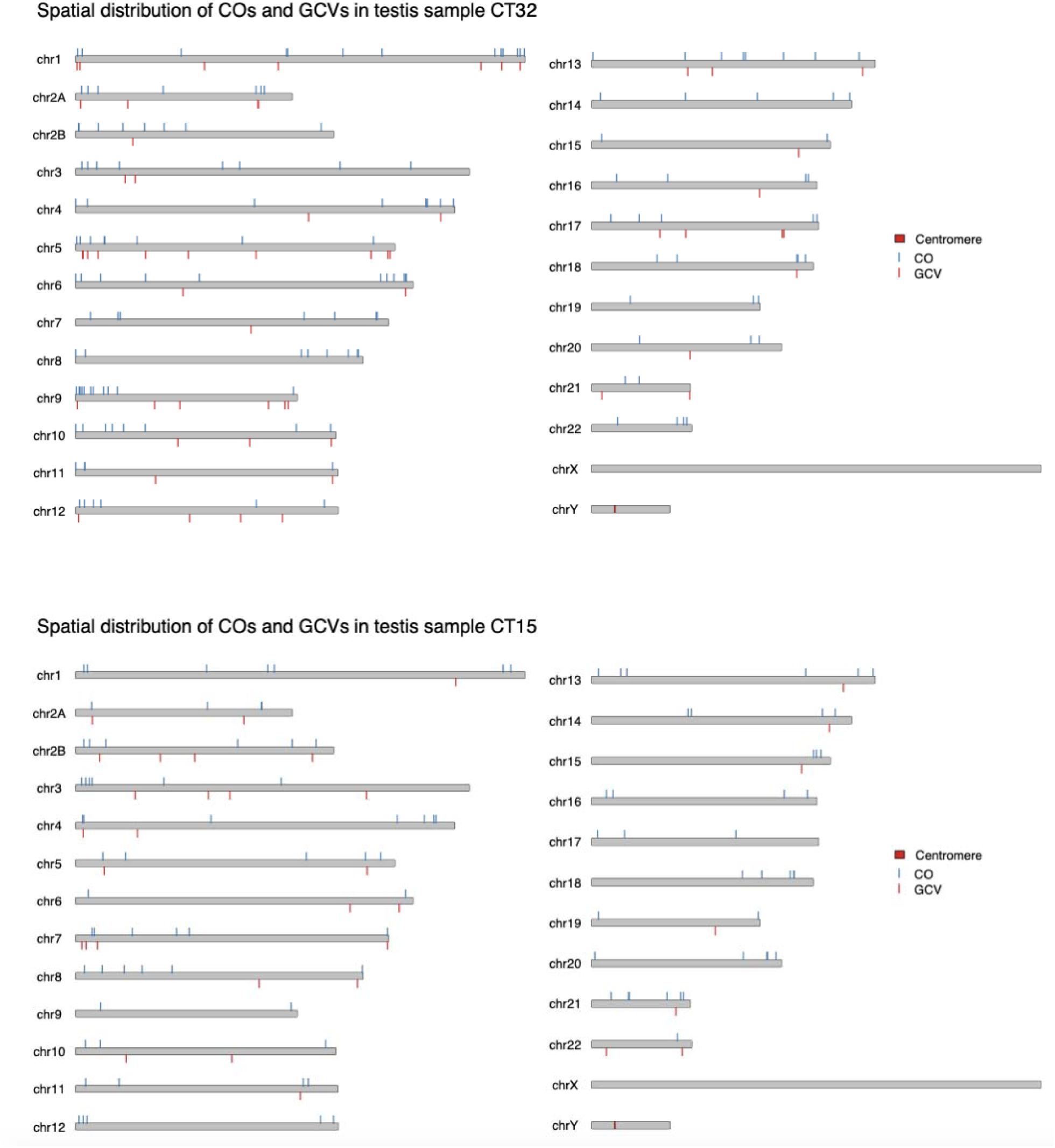
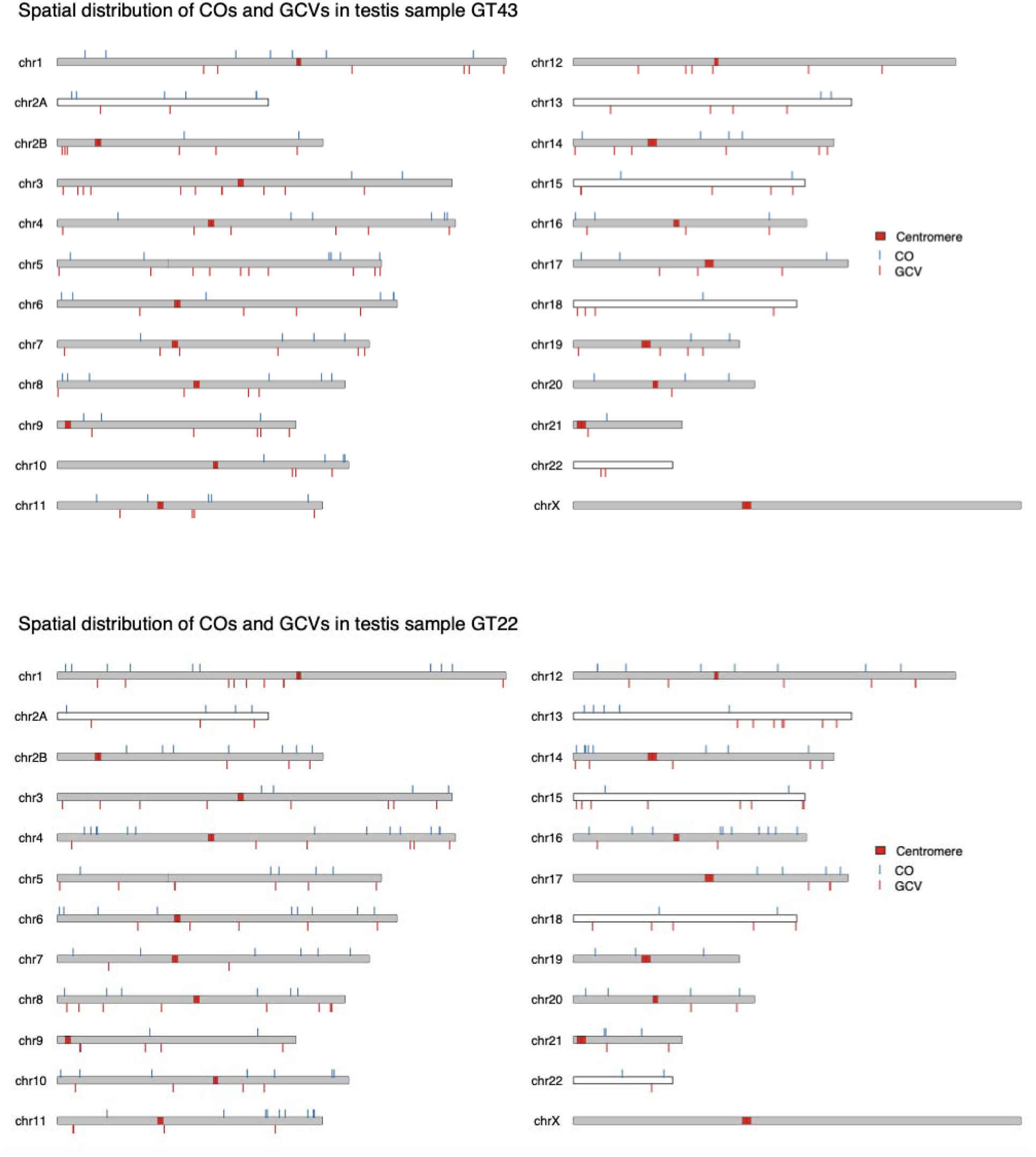
Chromosomal distribution of CO and GCV events for each of the samples Note: The X and Y chromosomes were assembled and represented here, no CO or GCV events were called on the X and Y chromosomes.

**Supplementary Figure 7.**
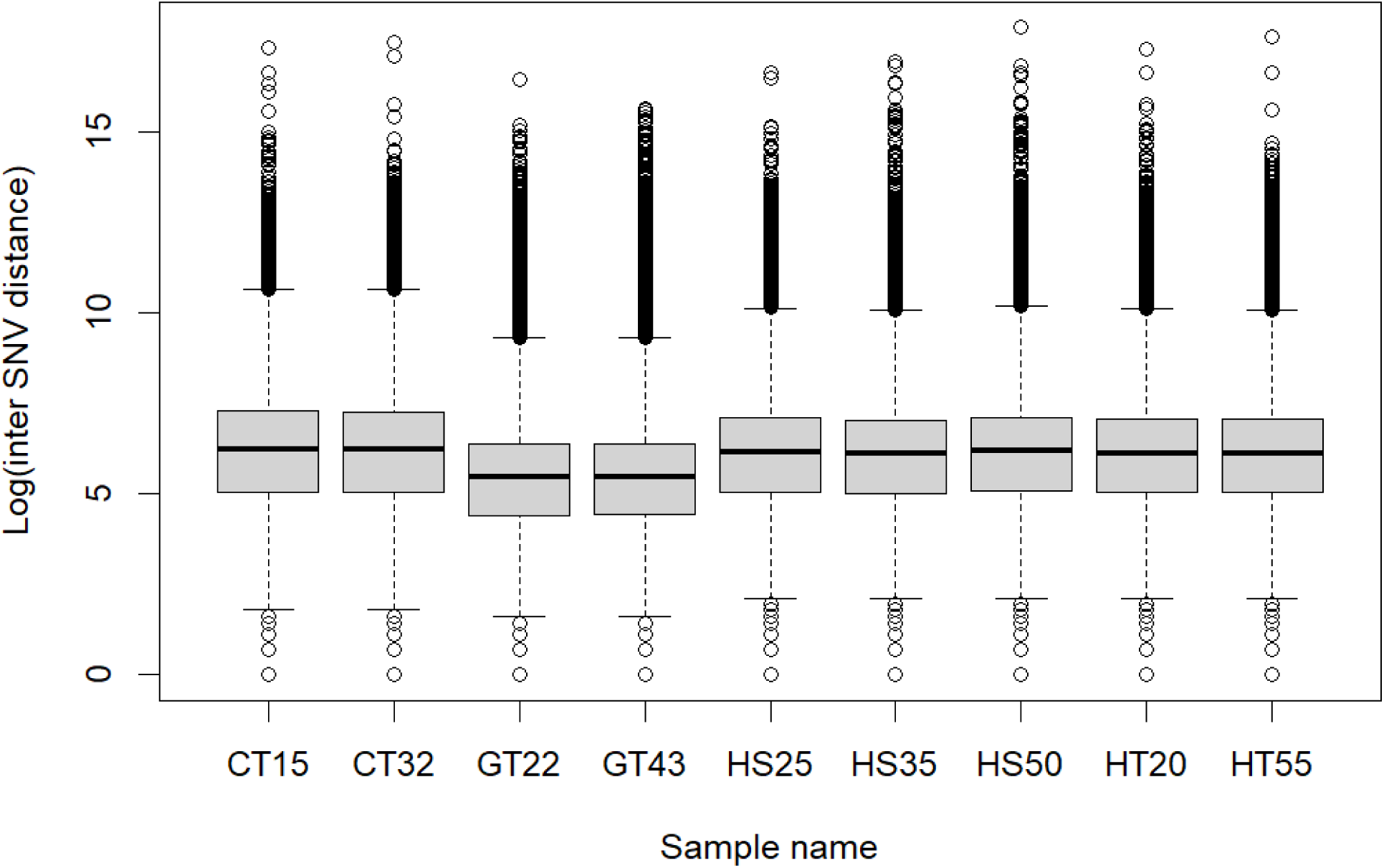
Boxplot showing the distribution of log inter-SNV distances throughout the genome of each sample. The results show that distances between SNVs vary by approximately 8 orders of magnitude throughout the genome, ranging from SNV clusters where SNVs are located next to each other to long runs of homozygosity where SNVs can be located up to ∼57 Mb from each other.

**Supplementary Table 1.** The table contains the average read length, the amount of the genome that is covered by contigs larger than 1 Mb, and the N50 for all nine samples. Individual identifiers end with each individual’s approximate age (eg 25, 35).

| Individual | Species (tissue) | Avg. read length (bp) | Contigs > 1 Mb (%) | N50 (Mb) |
| --- | --- | --- | --- | --- |
| HS25 | Human (sperm) | 16 362 | 97.9 | 70.7 |
| HS35 | Human (sperm) | 12 840 | 99.6 | 93.0 |
| HS50 | Human (sperm) | 17 148 | 99.4 | 89.4 |
| HT20 | Human (testis) | 14 302 | 98.0 | 92.4 |
| HT55 | Human (testis) | 10 616 | 97.6 | 74.7 |
| CT15 | Chimpanzee (testis) | 15 427 | 98.4 | 59.0 |
| CT32 | Chimpanzee (tests) | 15 118 | 93.3 | 72.7 |
| GT22 | Western Gorilla (testis) | 10 067 | 85.4 | 56.5 |
| GT43 | Western Gorilla (testis) | 11 015 | 92.9 | 50.8 |

**Supplementary Table 2:**
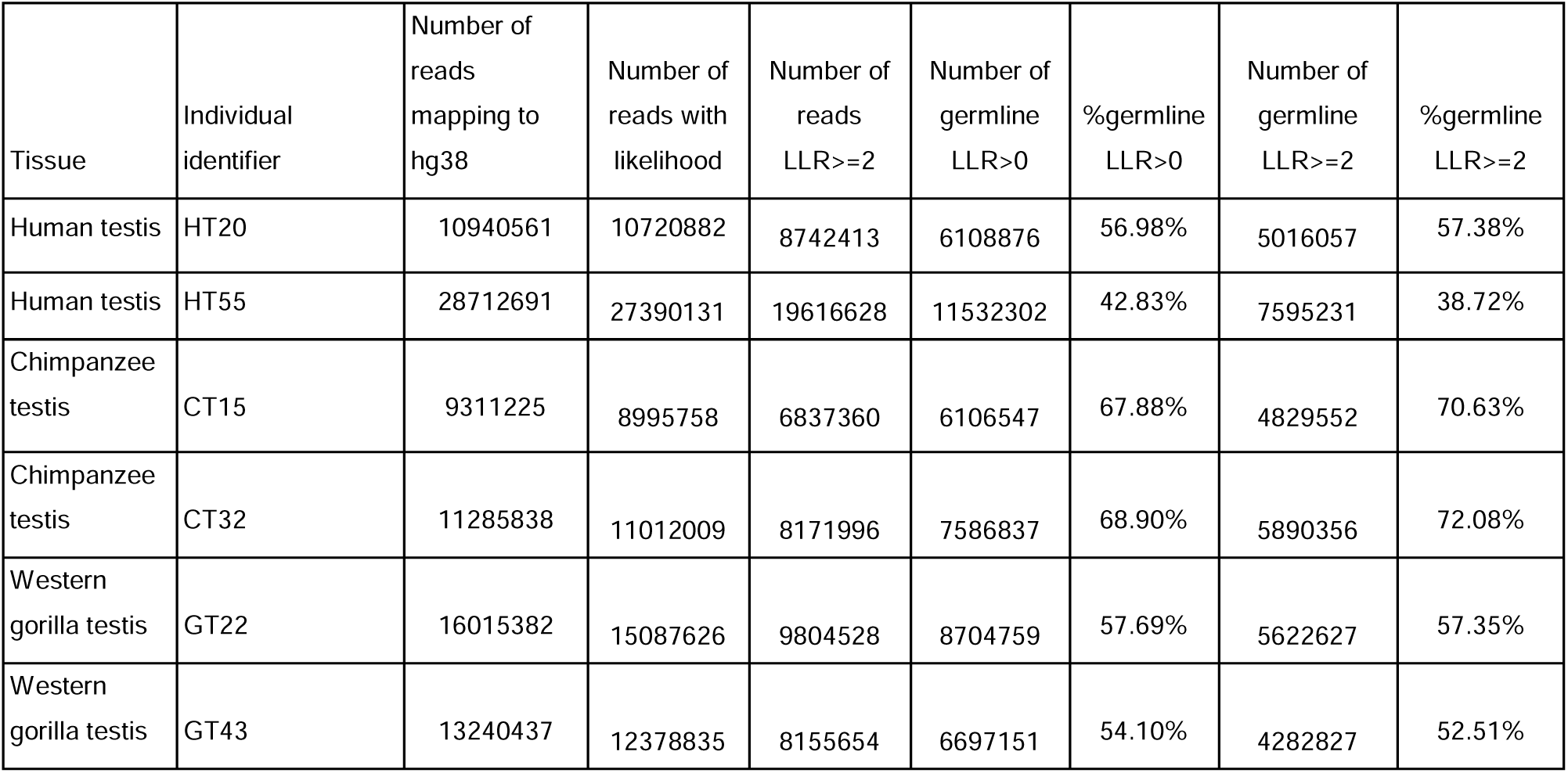
Summary of methylation-based classification of reads for each of the testis samples: Fraction of reads that can be classified with no cutoff (LLR>0) or by requesting a log-likelihood ratio of 2 or above (LLR>=2), and the fraction of reads classified with LLR>= 2 that are classified as germline for all reads and for each of the four recombination event types. Supplementary Table 3 shows the raw numbers that the fractions are based on.

| Tissue | Individual identifier | Number of reads mapping to hg38 | Number of reads with likelihood | Number of reads LLR>=2 | Number of germline LLR>0 | %germline LLR>0 | Number of germline LLR>=2 | %germline LLR>=2 |
| --- | --- | --- | --- | --- | --- | --- | --- | --- |
| Human testis | HT20 | 10940561 | 10720882 | 8742413 | 6108876 | 56.98% | 5016057 | 57.38% |
| Human testis | HT55 | 28712691 | 27390131 | 19616628 | 11532302 | 42.83% | 7595231 | 38.72% |
| Chimpanzee testis | CT15 | 9311225 | 8995758 | 6837360 | 6106547 | 67.88% | 4829552 | 70.63% |
| Chimpanzee testis | CT32 | 11285838 | 11012009 | 8171996 | 7586837 | 68.90% | 5890356 | 72.08% |
| Western gorilla testis | GT22 | 16015382 | 15087626 | 9804528 | 8704759 | 57.69% | 5622627 | 57.35% |
| Western gorilla testis | GT43 | 13240437 | 12378835 | 8155654 | 6697151 | 54.10% | 4282827 | 52.51% |

**Supplementary Table 3:**
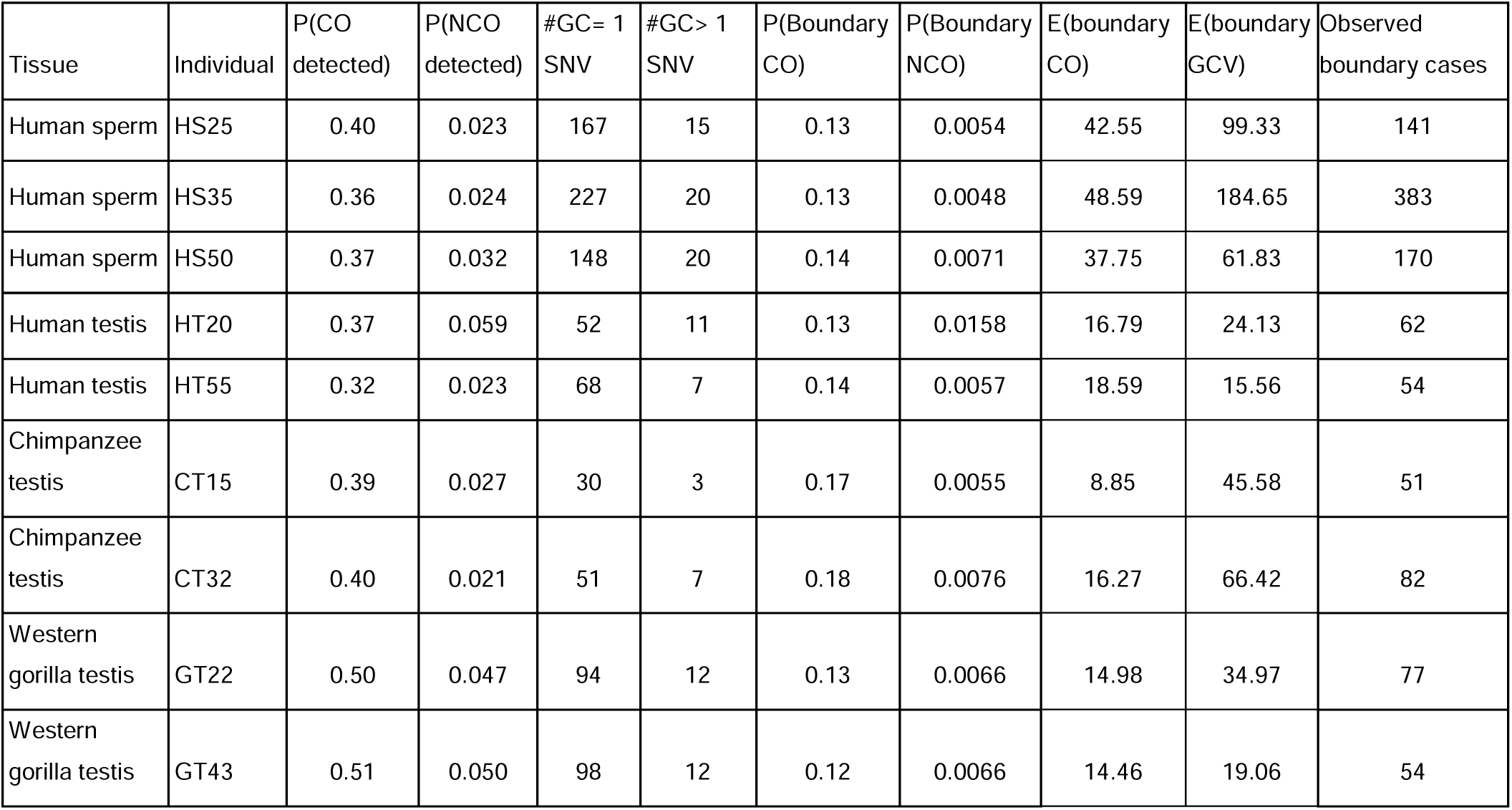
Detectability measures for crossovers (COs) and gene conversions (GCVs) estimated from simulations of events conditional on the read length distribution, the empirical SNV positions, and assuming that events are randomly distributed across the genome. #GC=1 and #GC>1 are the number of detected germline GCVs moving one SNV or more than one consecutive SNV, respectively. P(boundary CO) and P(boundary GCV) are the simulated probability that a CO and a GCV cause a shift in haplotype at the first or last SNV detected in a read. These are used to calculate how many of the boundary events are likely to be caused by COs and GCVs, respectively. The last column (from Table 1) is the observed number of boundary cases, which is close to the sum of the expected cases.

| Tissue | Individual | P(CO detected) | P(NCO detected) | #GC= 1 SNV | #GC> 1 SNV | P(Boundary CO) | P(Boundary NCO) | E(boundary CO) | E(boundary GCV) | Observed boundary cases |
| --- | --- | --- | --- | --- | --- | --- | --- | --- | --- | --- |
| Human sperm | HS25 | 0.40 | 0.023 | 167 | 15 | 0.13 | 0.0054 | 42.55 | 99.33 | 141 |
| Human sperm | HS35 | 0.36 | 0.024 | 227 | 20 | 0.13 | 0.0048 | 48.59 | 184.65 | 383 |
| Human sperm | HS50 | 0.37 | 0.032 | 148 | 20 | 0.14 | 0.0071 | 37.75 | 61.83 | 170 |
| Human testis | HT20 | 0.37 | 0.059 | 52 | 11 | 0.13 | 0.0158 | 16.79 | 24.13 | 62 |
| Human testis | HT55 | 0.32 | 0.023 | 68 | 7 | 0.14 | 0.0057 | 18.59 | 15.56 | 54 |
| Chimpanzee testis | CT15 | 0.39 | 0.027 | 30 | 3 | 0.17 | 0.0055 | 8.85 | 45.58 | 51 |
| Chimpanzee testis | CT32 | 0.40 | 0.021 | 51 | 7 | 0.18 | 0.0076 | 16.27 | 66.42 | 82 |
| Western gorilla testis | GT22 | 0.50 | 0.047 | 94 | 12 | 0.13 | 0.0066 | 14.98 | 34.97 | 77 |
| Western gorilla testis | GT43 | 0.51 | 0.050 | 98 | 12 | 0.12 | 0.0066 | 14.46 | 19.06 | 54 |

**Supplementary Table 4:** Estimates of NCO rates, NCO detection probability, NCO boundary case probability, and mean NCO tract length with CI95% intervals in brackets along with mean value estimates of CO detection probability and CO boundary case probability for all samples.

| Individual | NCO rate<br>[CI95%] | NCO detection<br>probability<br>[CI95%] | NCO boundary<br>probability<br>[CI95%] | CO boundary<br>probability<br>[CI95%] | CO detection<br>probability | Mean NCO<br>tract length<br>(bp) [CI95%] |
| --- | --- | --- | --- | --- | --- | --- |
| HS25 | 4.58e-06<br>[4.41e-06,<br>4.87e-06] | 0.0231<br>[0.0131,<br>0.0378] | 0.0054<br>[0.003,<br>0.0091] | 0.1336 | 0.4008 | 46<br>[25, 80] |
| HS35 | 2.83e-06<br>[2.72e-06,<br>3.01e-06] | 0.0244<br>[0.0157,<br>0.0381] | 0.0048<br>[0.003,<br>0.0079] | 0.1311 | 0.3621 | 47<br>[29, 78] |
| HS50 | 3.18e-06<br>[3.03e-06,<br>3.45e-06] | 0.0316<br>[0.0184,<br>0.0498] | 0.0071<br>[0.0044,<br>0.0117] | 0.1392 | 0.376 | 72<br>[40, 123] |
| HT20 | 3.82e-06<br>[3.38e-06,<br>4.59e-06] | 0.0593<br>[0.0311,<br>0.062] | 0.0158<br>[0.0078,<br>0.0191] | 0.1316 | 0.3654 | 140<br>[65, 280] |
| HT55 | 2.40e-06<br>[2.26e-06,<br>2.68e-06] | 0.023<br>[0.0092,<br>0.065] | 0.0057<br>[0.0022,<br>0.0197] | 0.1399 | 0.3237 | 45<br>[17, 105] |
| CT15 | 1.78e-06<br>[1.61e-06,<br>2.17e-06] | 0.0205<br>[0.0035,<br>0.0806] | 0.0055<br>[0.0009,<br>0.0114] | 0.1747 | 0.3948 | 45<br>[7, 175] |
| CT32 | 2.76e-06<br>[2.52e-06,<br>3.22e-06] | 0.0271<br>[0.0089,<br>0.0988] | 0.0076<br>[0.0024,<br>0.0294] | 0.1773 | 0.4004 | 60<br>[18, 160] |
| GT22 | 3.46e-06<br>[2.79e-06,<br>3.98e-06] | 0.0467<br>[0.0305,<br>0.0478] | 0.0066<br>[0.0043,<br>0.0129] | 0.1302 | 0.5027 | 55<br>[29, 95] |
| GT43 | 3.24e-06<br>[3.03e-06,<br>3.58e-06] | 0.0502<br>[0.0274,<br>0.0702] | 0.0066<br>[0.0035,<br>0.0106] | 0.1211 | 0.5084 | 45<br>[23, 80] |

**Supplementary Table 5:** The chromosome length in T2T coordinates, the length of segmental duplications (SD) and of the acrocentric regions that are masked for analysis. The last column is the callable fraction of each chromosome. Note: only chromosomes 13,14,15,21 and 22 have an acrocentric p-arm.

| Chromosome | Chromosome length (bases) | SD length (bases) | Acrocentric p-arm (bases) | Callable fraction (bases) |
| --- | --- | --- | --- | --- |
| Chr 1 | 248387328 | 15908974 | n.a | 232478354 |
| Chr 2 | 242696752 | 11658079 | n.a | 231038673 |
| Chr 3 | 201105948 | 4675018 | n.a | 196430930 |
| Chr 4 | 193574945 | 5304229 | n.a | 188270716 |
| Chr 5 | 182045439 | 5738023 | n.a | 176307416 |
| Chr 6 | 172126628 | 5738023 | n.a | 166388605 |
| Chr 7 | 160567428 | 13766937 | n.a | 146800491 |
| Chr 8 | 146259331 | 4312872 | n.a | 141946459 |
| Chr 9 | 150617247 | 13717227 | n.a | 136900020 |
| Chr 10 | 134758134 | 7765030 | n.a | 126993104 |
| Chr 11 | 135127769 | 5769583 | n.a | 129358186 |
| Chr 12 | 133324548 | 2464548 | n.a | 130860000 |
| Chr 13 | 113566686 | 3200481 | 15547593 | 94818612 |
| Chr 14 | 101161492 | 3191177 | 10092112 | 87878203 |
| Chr 15 | 99753195 | 8892475 | 16678794 | 74181926 |
| Chr 16 | 96330374 | 10851040 | n.a | 85479334 |
| Chr 17 | 84276897 | 8262691 | n.a | 76014206 |
| Chr 18 | 80542538 | 2305844 | n.a | 78236694 |
| Chr 19 | 61707364 | 4367829 | n.a | 57339535 |
| Chr 20 | 66210255 | 3282089 | n.a | 62928166 |
| Chr 21 | 45090682 | 1573782 | 10962853 | 32554047 |
| Chr 22 | 51324926 | 7497258 | 12788180 | 31039488 |

**Supplementary Table 6:** The number of called events and the number of events that passes the manual curation in all four categories.

| Individual | Called CO events | Approved CO events | Called GCV events | Approved GCV events | Called Boundary events | Approved Boundary events | Called Complex events | Approved Complex events |
| --- | --- | --- | --- | --- | --- | --- | --- | --- |
| HS25 | 320 | 298 | 200 | 182 | 241 | 141 | 41 | 34 |
| HS35 | 566 | 510 | 279 | 247 | 538 | 383 | 27 | 10 |
| HS50 | 352 | 321 | 190 | 168 | 254 | 170 | 40 | 24 |
| HT20 | 84 | 67 | 82 | 63 | 117 | 62 | 15 | 10 |
| HT55 | 72 | 36 | 90 | 74 | 239 | 54 | 27 | 19 |
| CT15 | 125 | 100 | 59 | 33 | 106 | 51 | 34 | 11 |
| CT32 | 183 | 149 | 87 | 57 | 154 | 82 | 26 | 14 |
| GT22 | 162 | 132 | 160 | 102 | 149 | 77 | 40 | 18 |
| GT43 | 105 | 79 | 146 | 106 | 113 | 54 | 46 | 13 |

